# The Fate of Glutamine in Human Metabolism. The interplay with glucose in proliferating cells

**DOI:** 10.1101/477224

**Authors:** Jean-Pierre Mazat, Stéphane Ransac

**Affiliations:** IBGC CNRS UMR 5095 & Université de Bordeaux, 1, rue Camille Saint-Saëns 33077 BORDEAUX-cedex, France

**Keywords:** Model of central carbon and nitrogen metabolism, Flux Balance Analysis, Glutamine Warburg effect, hypoxia

## Abstract

Genome-scale models of metabolism (GEM) are used to study how metabolism varies in different physiological conditions. However, the great number of reactions involved in GEM makes it difficult to understand these variations. In order to have a more understandable tool, we developed a reduced metabolic model of central carbon and nitrogen metabolism, C2M2N with 77 reactions, 54 internal metabolites and 3 compartments, taking into account the actual stoichiometry of the reactions, including the stoichiometric role of the cofactors and the irreversibility of some reactions. In order to model OXPHOS functioning, the proton gradient through the inner mitochondrial membrane is represented by two pseudo-metabolites DPH (ΔpH) and DPSI (Δψ).

To illustrate the interest of such a reduced and quantitative model of metabolism in mammalian cells, we used Flux Balance Analysis (FBA), to study all the possible fates of glutamine in metabolism. Our analysis shows that glutamine can supply carbon sources for cell energy production and can be used as carbon and nitrogen sources to synthesize essential metabolites. Finally, we studied the interplay between glucose and glutamine for the formation of cell biomass according to ammonia microenvironment. We then propose a quantitative analysis of the Warburg effect.

## 1. Introduction

Genome-scale models of metabolism (GEM) greatly help to understand how metabolism varies in different physiological conditions, in different environments, in case of enzyme deficiencies and in interaction with other metabolisms. Used in conjunction with methods such as flux balance analysis [1], GEM are particularly useful to simulate metabolic changes in large metabolic networks as they do not require kinetic parameters and are computationally inexpensive. Many genome-scale constraint-based models [2–8] have covered central metabolism and have been used successfully to model diseases [9,10]. However, the great number of reactions involved in GEM makes it difficult to understand the results obtained in these kinds of studies. In order to have a more manageable tool, we developed a reduced metabolic model of central carbon and nitrogen metabolism, C2M2N, taking into account the true stoichiometry of the reactions, including the stoichiometric role of the cofactors and the irreversibility of some reactions. The configuration used in this work involves three compartments: the extracellular medium, the cytosol and the mitochondrial matrix. 77 reactions among 37 are irreversible and 54 internal metabolites are taken into account. Mitochondrial metabolism (OXPHOS) and mitochondrial transports, are often inaccurately represented in GEM with some exceptions [2,11]. In C2M2N, the model of OXPHOS functioning is based on a proton gradient through the inner mitochondrial membrane, represented here by two pseudo-metabolites DPH (ΔpH) and DPSI (Δψ). This allowed us to specifically take into account the ‘vectorial’ protons across the mitochondrial membrane and not the protons in the reactions inside a given compartment which are, in principle, equilibrated. The main purpose in developing our core model was to create a metabolic model as small as possible, involving the main reactions encountered in experimental metabolic studies to be studied with several theoretical approaches, including the determination of Elementary Flux Modes (EFMs) [13].

To illustrate the practical interest of such a reduced model of metabolism in mammalian cells, we studied glutamine’s metabolism and compared it with glucose metabolism. Glutamine is the most abundant amino acid in plasma and has long been recognized to be essential in proliferating cells. Additionally, glutamine was identified as an alternative to glucose to fuel the Krebs cycle in cancer cells or in hypoxia conditions or mutations [14–20]. The glutamine metabolism goes through glutamate and α-ketoglutarate (AKG). Glutamate can be produced in the mitochondria by glutaminase or transaminases and in the cytosol through nucleotide synthesis or transaminases. When synthesized inside the mitochondria, glutamate exits through the H^+^/Glutamate co-transporter [21] (the glutamate/aspartate exchanger with a H^+^ entry is sensitive to the ΔµH^+^and can be considered as irreversible in normal physiological conditions). Inside the mitochondria, glutamine-derived AKG replenishes the TCA cycle and can be metabolized either through the canonical forward mode or via reductive carboxylation, leading to citrate and acetyl-CoA in the cytosol. For instance, Chen et al. [22] showed that glutamine enables the survival of mitochondrial DNA mutant cells thanks to both reductive and oxidative pathway in the TCA cycle.

Using Flux Balance Analysis (FBA), we systematically studied and quantitatively discussed all the possible fates of glutamine in central carbon metabolism and demonstrated that glutamine can sustain cell energy production and be used as a carbon and nitrogen source to synthesize essential metabolites, thus contributing to cell proliferation. Here, we show how C2M2N can be used to explore the results of more complex metabolic models by comparing our results with those of MitoCore, a metabolic model of intermediate size built on the same bases.

In a more general approach, we used C2M2N to follow how glucose and glutamine together share energy metabolism and anabolism in different physiological or pathological conditions (proliferating cells, hypoxia). We demonstrated the interest of a core stoichiometric model such as C2M2N in quantitatively tracing the origin of the carbon and nitrogen atoms in the different syntheses and in specifying the respective role of glutamine and glucose in energy metabolism. We demonstrated the role of ammonia release or input or recycling and study some Urea Cycle Dysregulation (UCD). Ultimately, we gave a quantitative account of Warburg effect evidencing the main challenge of proliferating cells, i.e. energy production while maintaining NAD/NADH balance.

## 2. Short description of the metabolic models

### 2.1. C2M2N Model

The reactions involved in the metabolic model C2M2N (Central Carbon Metabolic Model with Nitrogen) are listed in Appendix B and depicted in Figure 1. C2M2N was developed by trial and error from the traditional metabolic networks that can be found in textbooks with the aim to get the smallest network that accounts for most metabolic functions. To this aim, aggregation of consecutive reactions was extensively used when possible while keeping the relevant stoichiometry, particularly of the cofactors. However, C2M2N is not strictly defined, new reactions can be added when necessary (see reaction DH below) or easily suppressed by constraining their flux to zero. The abbreviations of the reaction names are listed in Appendix A. More precisely, the model consists in (i) a simple version of the Krebs cycle and connected reactions (reactions PDH, CS, IDH2 and 3, SLP, RC2, MDH2 and PYC with the addition of glutamate dehydrogenase (GLUD1)), (ii) in the glycolysis summarized in 5 steps, G1 (hexokinase + phosphoglucose isomerase), G2 (phosphofructokinase + aldolase + triose-phosphate isomerase), G3 (glyceraldehyde-3P dehydrogenase + phosphoglycerate kinase), ENOMUT (enolase + phosphoglycerate mutase) and PK (pyruvate kinase) extended by the reversible LDH (lactate dehydrogenase) and the possible output/input of lactate (LACIO). Gluconeogenesis consists in the reversible reactions of glycolysis plus PEPCK1 (phosphenolpyruvate carboxykinase), GG3 (triose phosphate isomerase + aldolase + fructose-1,6-biphosphatase) and GG4 (phosphogluco isomerase + glucose-6-phosphatase). The mitochondrial phosphenolpyruvate carboxykinase named PEPCK2 was included. The pentose phosphate pathway (PPP) reactions are summarized in PP1 (oxidative part of PPP) and PP2 (non-oxidative part of PPP). The urea cycle is summarized in 4 reactions: CPS1_OTC (carbamoyl phosphate synthase 1 + Ornithine transcarbamylase), ORNT1 (Ornithine/Citrulline + H^+^ exchanger), ASS1_ASL_FH (argininosuccinate synthase + argininosuccinate lyase to which we added the fumarate hydratase to convert fumarate in malate as we did for RC2 reaction) and finally ARGASE (arginase). The ornithine synthesis from glutamate and acetylCoA, NAGS_ACY (N-acetylglutamate synthase + Amino Acylase) is completed by the regeneration of acetyl_CoA from acetate by ACS (AcetylCoA Syntethase) and by the ornithine/H^+^ transporter ORNT2. The urea cycle is supplemented by NOS (NO Synthase).

**Figure 1.**
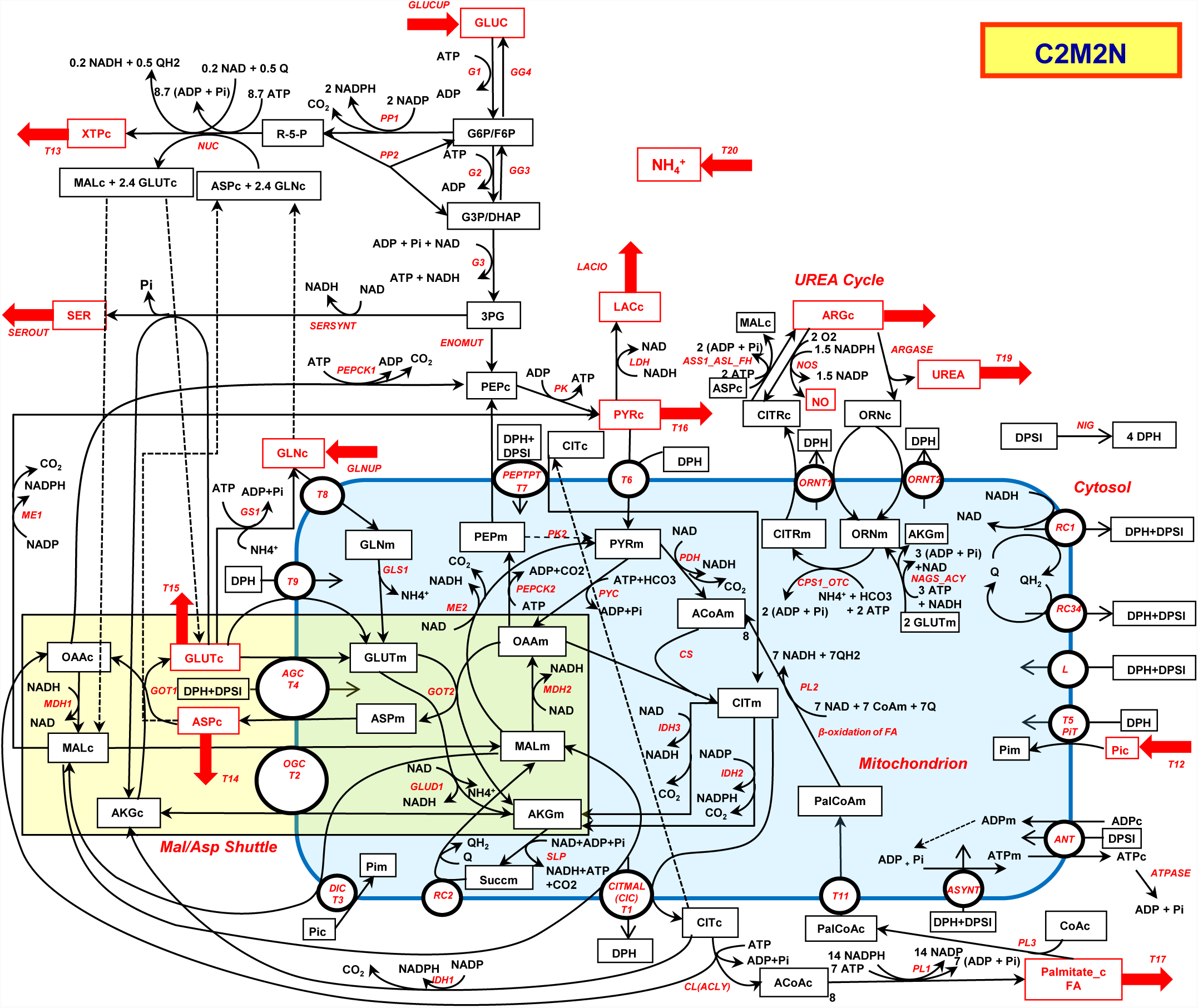
C2M2N Central Carbon Metabolic with Nitrogen Model. Possible entries and outputs are indicated by red arrows. The corresponding metabolites are in red. The abbreviated names of the reactions are indicated in red italic along the arrows of the reactions. The mitochondrion is in blue and the malate/aspartate shuttle in light brown. Dotted lines link identical metabolites duplicated for the sake of presentation.

The synthesis of nucleotide bases (NUC) is represented by a simplification of purine and pyrimidine biosynthesis obtained by averaging the stoichiometry of the different metabolites and cofactors in the metabolic pathway of each nucleotide and taking into account their different amounts in human (30% of A and T or U and 20% of G and C). It should be stressed that nucleotide synthesis necessitates glutamine which is converted to glutamate and aspartate, itself converted to fumarate and subsequently to malate.

The synthesis of serine from 3-phosphoglycerate involves 3 steps: a dehydrogenase, a transaminase involving the glutamate/ α-ketoglutarate pair and a phosphatase. These three steps are assembled in one reaction: SERSYNT.

The malate/aspartate shuttle (MAS) is fully represented in the direction of NADHc consumption and NADHm production. It involves the malate/ α-ketoglutarate exchanger T2 (OGC) and the glutamate/aspartate exchanger T4 (AGC), the malate dehydrogenases (cytosolic MDH1 and mitochondrial MDH2) and the glutamate-oxaloacetate transaminases (cytosolic GOT1 and mitochondrial GOT2). A detailed representation of MAS was mandatory because MAS enzymes are not always operating with the stoichiometry and the direction of MAS for exchange of NADHc for NADHm i.e. MAS components are not always used to run the MAS as such.

The synthesis of fatty acids is a major pathway in proliferating cells. It starts with citrate lyase (CL) and is represented in the case of palmitate by the reaction PL1 with the corresponding stoichiometries.

The respiratory chain is represented by three reactions, RC1 which is the respiratory complex I, RC2 (succinate dehydrogenase or complex II which also belongs to the TCA cycle + fumarase) and RC34 which represents complex III + IV.

Finally, the remaining steps of oxidative phosphorylation are represented by ASYNT (ATP synthase), ANT, the ADP/ATP exchanger, T5 the Pi carrier and L the protons membrane leak. The proton gradient is represented by two pseudo-metabolites DPH (ΔpH) and DPSI (Δψ). It allowed us to only take into account the ‘vectorial’ protons across the mitochondrial membrane [12]. We introduced a possible ΔpH / Δψ exchange (NIG), mimicking the action of nigericin (K^+^/H^+^ exchanger) or physiologically, the action of ion exchangers in the inner mitochondrial membrane. DPH and DPSI are introduced, when necessary, in mitochondrial carriers equations. ATP hydrolysis is symbolized by ATPASE activity.

Only two entries are considered here, the entry of glutamine (GLNUP) and the entry of glucose (GLUCUP). The possible outputs in this study are pyruvate (T16), serine (SEROUT), aspartate (T14) nucleotides (T13) and palmitate (T17), arginine (T18) urea (T19) and ammonia (T20). The entry of glutamine in the mitochondria goes through the transporter T8. It will give rise to glutamate inside the mitochondria through glutaminase GLS1.

### 2.2. MitoCore Model

MitoCore [11] is a manually curated constraint-based computer model of human metabolism that incorporates 324 metabolic reactions, 83 transport steps between mitochondrion and cytosol, and 74 metabolite inputs and outputs through the plasma membrane, to produce a model of manageable scale for an easier interpretation of results. The representation of the proton gradient is nearly the same as in C2M2N with the pseudo substrates PMF (Proton Motive Force) with DPH = 0.18 PMF and DPSI = 0.82 PMF. MitoCore’s default parameters simulate normal cardiomyocyte metabolism with entry of different metabolites, particularly glucose and amino acids. These entries were lowered to zero and glutamine or glucose entry were set to one (unless otherwise mentioned) to compare the results of MitoCore with those of C2M2N. (See the SBML file of MitoCore used in this study in the supplementary materials).

### 2.3. FBA analysis

We used FAME (Flux Analysis and Modeling Environment) (http://f-a-m-e.org/) [23] to derive the set of fluxes of C2M2N (and of MitoCore for comparison) optimizing the objective functions used below with glutamine or glucose entry for the synthesis of various metabolites. We systematically looked for the absolute flux minimization and then used the Flux Variability Analysis (FVA) to get an idea of the possible other solutions giving the same objective functions. Among these solutions (with a maximal objective function), we selected and drew the one with the maximal ATP synthesis rate. In the main text, the solutions on a simplified version of Figure 1 were represented (Figure 2 and the following). The complete representations of the C2M2N fluxes are in the supplementary materials with the corresponding figures labelled Figure Si for the simplified Figure i. The fates of glucose carbons are represented in blue and the fates of glutamine carbons are represented in green. When syntheses arise from both glutamine and glucose other colours are used. Results are given in Table 1.

**Table 1.** Maximal yield in metabolites synthesis or energy (ATP) from glutamine and glucose at steady state. The first value is obtained with C2M2N and the second with MitoCore.

| Objective Function | 1 x GLN |  | 1 x GLUC |  |
| --- | --- | --- | --- | --- |
|  | Metabolite | ATP | Metabolite | ATP |
| ATPASE | - | 23.8 / 24.0 | - | 33.34 / 33.04 |
| Glutamate (T15) | 1 / 1 | 0 / 0 | 1 | 8.9 / 8.5 |
| Pyruvate (T16) | 1 / 1 | 10.6 / 10.9 | 2 / 2 | 6.9 / 6.85 |
| Aspartate (T14) | 1 / 1 | 7.6 / 7.7 | 1.85 / 1.82 | 2 / 2 |
| XTP (T13) | 0.6 / - | 1 / - | 0.86 / - | 0 / - |
| Palmitate (T17) | 0.17 / 0.17 | 0 / 0 | 0.25 / 0.24 | 0 / 0 |
| Serine (SEROUT) | 1 / 1 | 9.3 / 9.1 | 2 / 2 | 4.4 / 3.8 |

**Figure 2.**
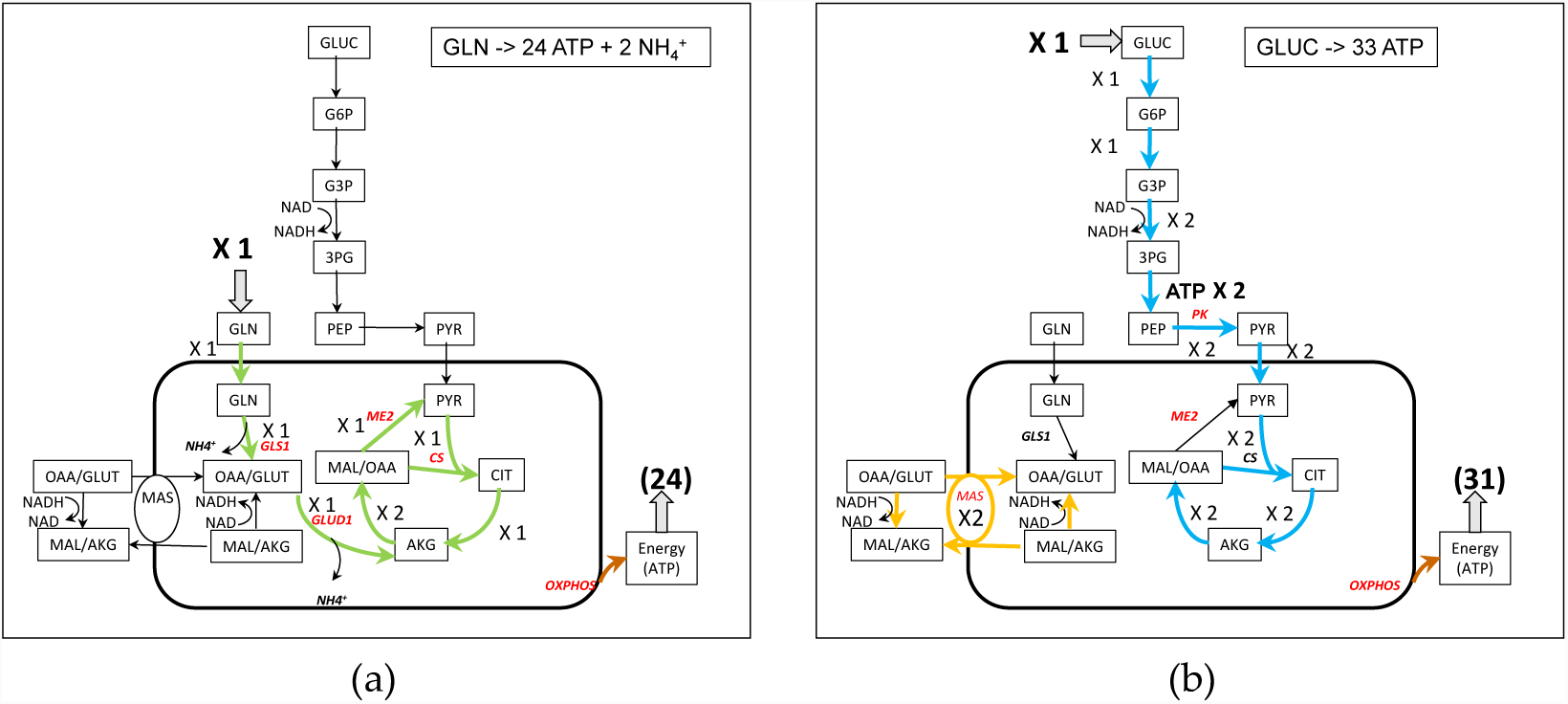
Simplified representation of ATP synthesis from glutamine (a) and glucose (b). Note that in (b), for the sake of simplicity, the representation of MAS has been separated from the TCA cycle although they share the MDH2 activity with a net flux of 4, 2 for MAS and 2 for the TCA cycle. Complete figures of the fluxes in Figure S2.

### 2.4. Optimization of proliferating cells biomass

In order to understand how glutamine and glucose share the increase in Biomass of proliferating cells, we maximized a Biomass function (BM) defined from [24]:

BM = 0.18 SERc + 0.11 Palmitate_c + 0.04 XTPc + 0.19 GLNc + 0.17 GLUTc + 0.7 ASPc + 0.05 ARGc + 0.12 PYRc + 4 ATPc = => 1 Biomass + 4 ADPc + 4 Pic.

PYRc takes the place of alanine, ARG summarizes arginine and proline in biomass and also polyamines synthesis, and SER is for serine + glycine.

## 3. Results and discussions

### 3.1. Use of Glutamine for energy production (Figure 2)

Energy production is symbolized by ATPASE activity which is the objective function in this section. About 24 molecules of ATP can be synthesized from 1 molecule of glutamine without any other synthesis (Figure 2a and S2a). One molecule of glutamine enters the mitochondria and then the Krebs cycle as α-ketoglutarate (AKG) to double the flux to malate. One molecule of malate generates pyruvate (through ME2) and then acetyl-CoA which will condense with the OAA derived from the remaining molecule of malate to generate citrate and a canonical TCA cycle continue in the usual direction with a flux equal to 1. It is very similar to the glycolysis-derived pyruvate TCA cycle except that, with glutamine, there is no need for NADHc reoxidation. However, ATP synthesis from glutamine necessitates the synthesis of pyruvate either inside or outside mitochondria (see the discussion in [25,26] and is accompanied by a release of ammonia. In the same conditions, one molecule of glucose usually allows the synthesis of 33 ATP molecules at the most (Figure 2b and S2b). In both cases, the activity of respiratory chain is mandatory in order to reoxidise the NADHm produced by the TCA cycle and to get the maximum rate of ATP synthesis.

### 3.2. Pyruvate synthesis from Glutamine (Figure 3)

The production of pyruvate (precursor of alanine) from glutamine can follow the production of glutamate and α-ketoglutarate (AKG) entering the “left”, -oxidative-part of TCA cycle to produce OAA. From OAA, mitochondrial PEPCK2 produces PEP, which exits the mitochondria through citrate and malate cycling (grey arrows on Figure S3a) and generates pyruvate (with PK). One molecule of pyruvate is obtained per molecule of glutamine and 10.6 molecules of ATP can be synthesized in this condition. Another solution without ATP synthesis is depicted in Figure S3c and the more complex solution with MitoCore in Figure S3d. These solutions are analysed in supplementary materials.

**Figure 3.**
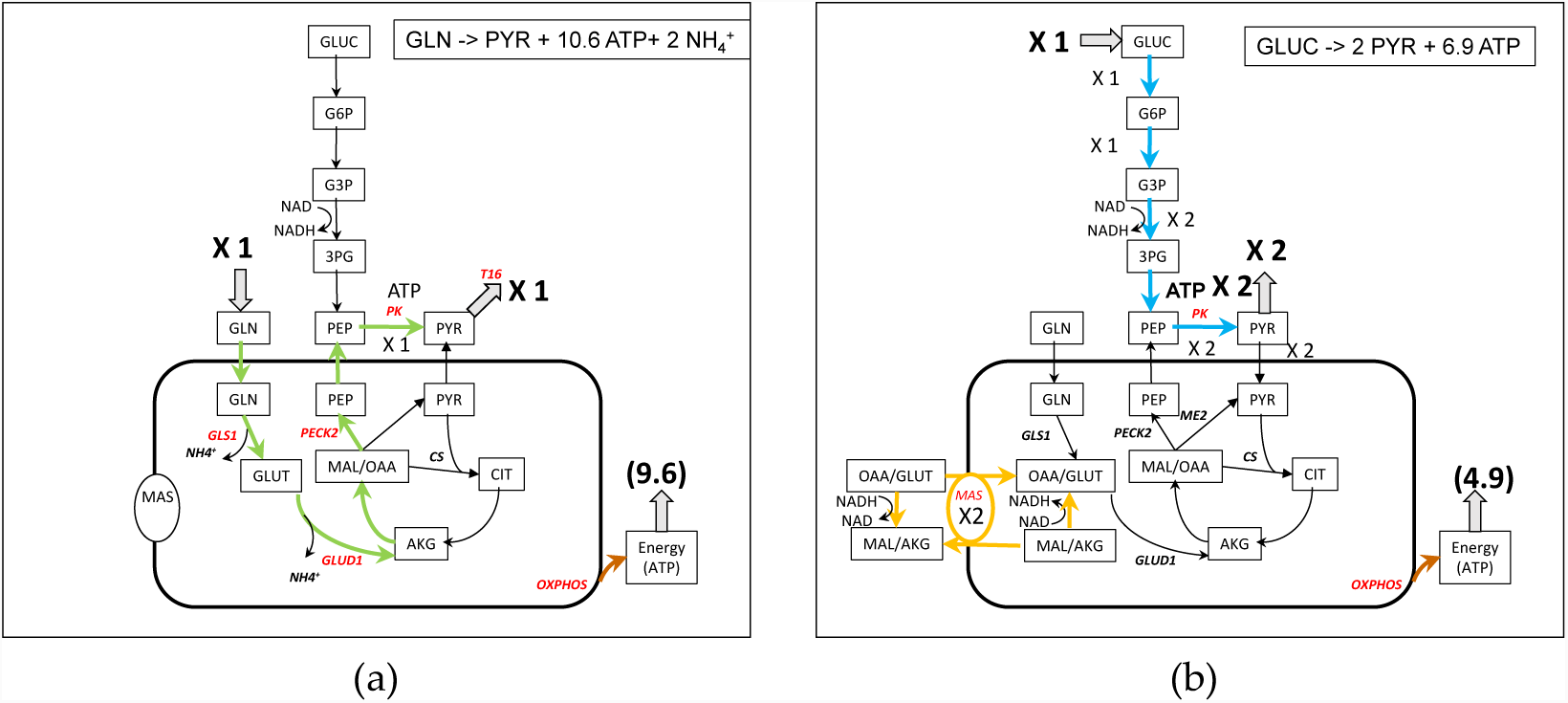
Simplified representation of pyruvate synthesis from glutamine (a) and glucose (b). Note that in (b), the glycolytic flux and the MAS flux are linked by NAD/NADH cycling. Complete figures of the fluxes in Figure S3.

With glucose as substrate (Figure 3b), the solution is rather simpler, with the synthesis of 2 glycolytic pyruvate and the reoxidation of NADHc by MAS and ETC accompanied by the corresponding ATP synthesis (6.9).

### 3.3 Apartate biosynthesis from glutamine (Figure 4)

Aspartate is an amino acid which participates in many reactions, particularly in nucleotides synthesis and appears at a high level in the biomass. Due to its low concentration in blood, aspartate synthesis is crucial for cell survival [27–31]. Aspartate synthesis from glutamine (Figure 4a and S4a) involves aspartate amino transferase (GOT) with oxalacetate (OAA) synthesized in the last part of TCA cycle from AKG coming from the glutamine-derived glutamate. Aspartate is carried out of the mitochondria through the glutamate-aspartate carrier (T4). Glutamate is recycled outside the mitochondria by the glutamate/H^+^ carrier (T9). The maximal yield from one glutamine is one aspartate and 7.6 ATP.

**Figure 4.**
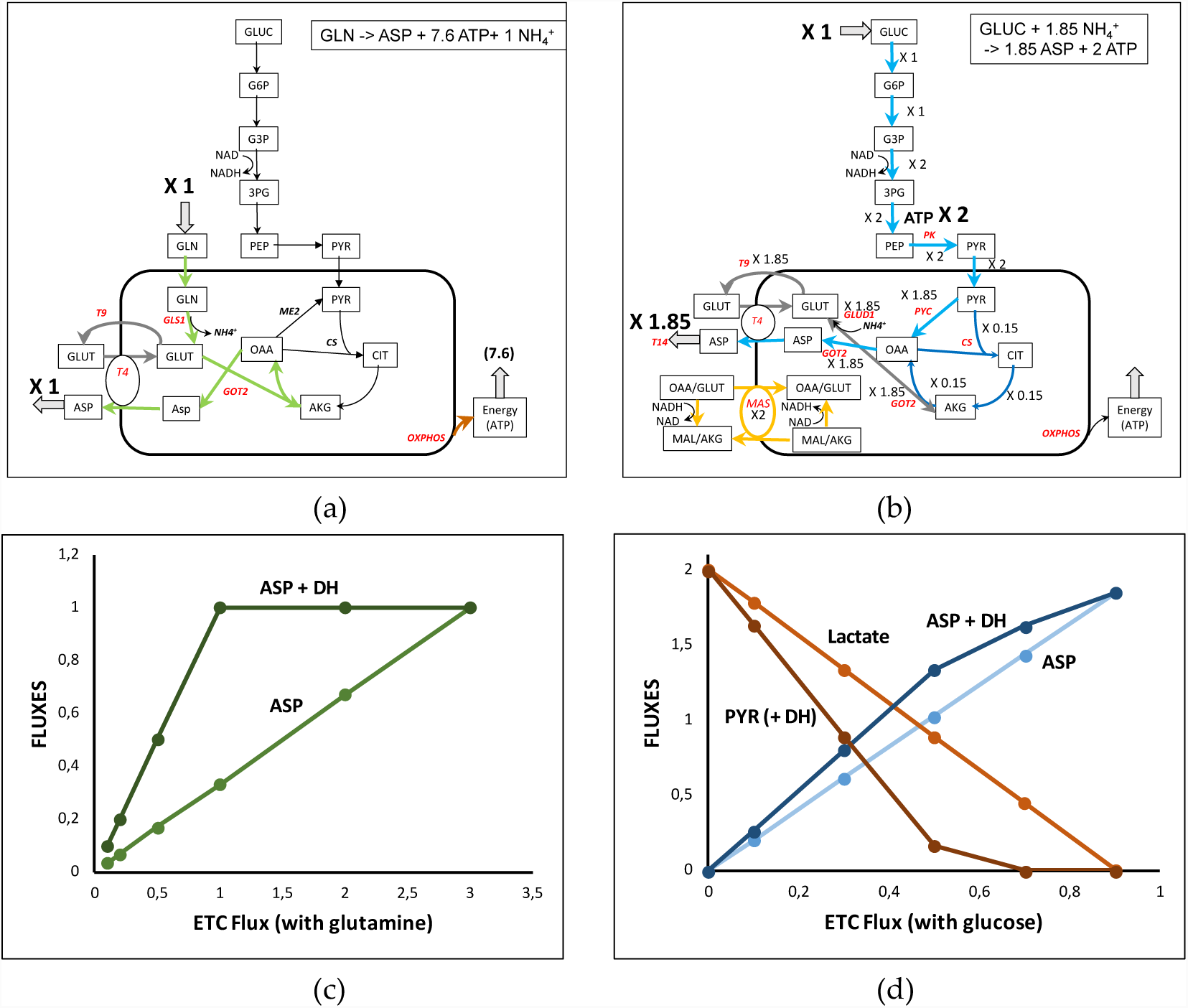
Simplified representation of aspartate synthesis from glutamine (a) and glucose (b). Note that in (b), the flux through T4 and GOT 2 can be split in a malate-aspartate shuttle (MAS) with a flux of 2 and in a 1.85 flux through T4 corresponding to aspartate output, i.e. T4 net flux is 3.85. See the complete representation of the glutamine- and glucose-derived aspartate in Figure S4a and b. (c): ETC inhibition (RC34 is inhibited) of aspartate synthesis from glutamine in the absence or presence of the dehydrogenase DH which reoxidizes NADHc. (d) ETC inhibition (RC34 is inhibited) of aspartate synthesis from glucose in the absence or presence of the dehydrogenase DH which reoxidizes NADHc. The ETC inhibition leads to lactate release, or pyruvate release in the presence of DH.

Aspartate synthesis from glucose also requires the activity of GOTs (cytosolic or mitochondrial) but is more complicated (Figure 4b and in its full representation in Figure S4b). In the solution of Figure 4b and S4b, 1.85 aspartate molecules and 2 glycolytic ATP are synthesized per glucose. Reoxidation of glycolytic NADH imposes the operation of the malate-aspartate shuttle (MAS). NADH cannot be reoxidized by the lactate dehydrogenase because pyruvate carbons are necessary for aspartate synthesis. For this reason, in Figure 4b MAS is represented with a flux equal to 2 and the glutamate-aspartate carrier (T4) with a supplementary activity of 1.85 to release the 1.85 aspartate outside the mitochondria. More precisely on Figure S4b one can see that the flux of 2 pyruvate entering mitochondria is split in 1.85 pyruvate carboxylase flux and a 0.15 canonical TCA cycle (thin dark blue arrows in Fig. 4b and S4b) generating the 1.85 ATPm necessary for the operation of 1.85 pyruvate carboxylase.

In recent papers [27–29] some authors evidenced “an essential role of the mitochondrial electron transport chain… in aspartate synthesis”. This is not unexpected if we consider that mitochondrial synthesis of aspartate requires the synthesis of OAAm (using pyruvate carboxylase with ATP or malate dehydrogenase generating NADH) and the fact that the GLU/ASP exchanger (T4) depends on the ΔµH^+^, i.e. the ETC activity. We can confirm this result in our model as represented in Figure 4c and d, which depicts the synthesis of aspartate as a function of ETC activity with one glutamine or one glucose. In both cases, the flux of aspartate synthesis proportionally decreases with the decrease of the activity of respiratory chain (modulated by RC34 activity). As shown experimentally, this is due in part to the necessity to reoxidize NADHc. Introducing in C2M2N a dehydrogenase activity (DH) reoxidizing NADHc, we induce an increase of aspartate synthesis in accordance with the results in [29].

Other reactions also appear essential such as the pyruvate carboxylase (PYC) and the pentose phosphate pathway.

### 3.4 Nucleotide synthesis from glutamine and glucose (Figure 5)

The synthesis of nucleotides (XTP) requires glutamine, aspartate and R5P. In the absence of ASP and R5P, glutamine has to be used to ensure these syntheses. In the solution of figure 5a, 0.6 XTP is synthesized that necessitates 1.44 glutamine. To do so, the input of one glutamine is used with, in addition, a 0.44 cycling of glutamate/glutamine via glutamine synthase (GS1). The glutamate resulting from glutamine entry (flux = 1) is entirely transformed in PEP then R5P through gluconeogenesis and pentose phosphate pathway. This solution necessitates a slight entry of NH_4^+^_ (0.04) and is accompanied by the production of 0.98 ATP.

**Figure 5.**
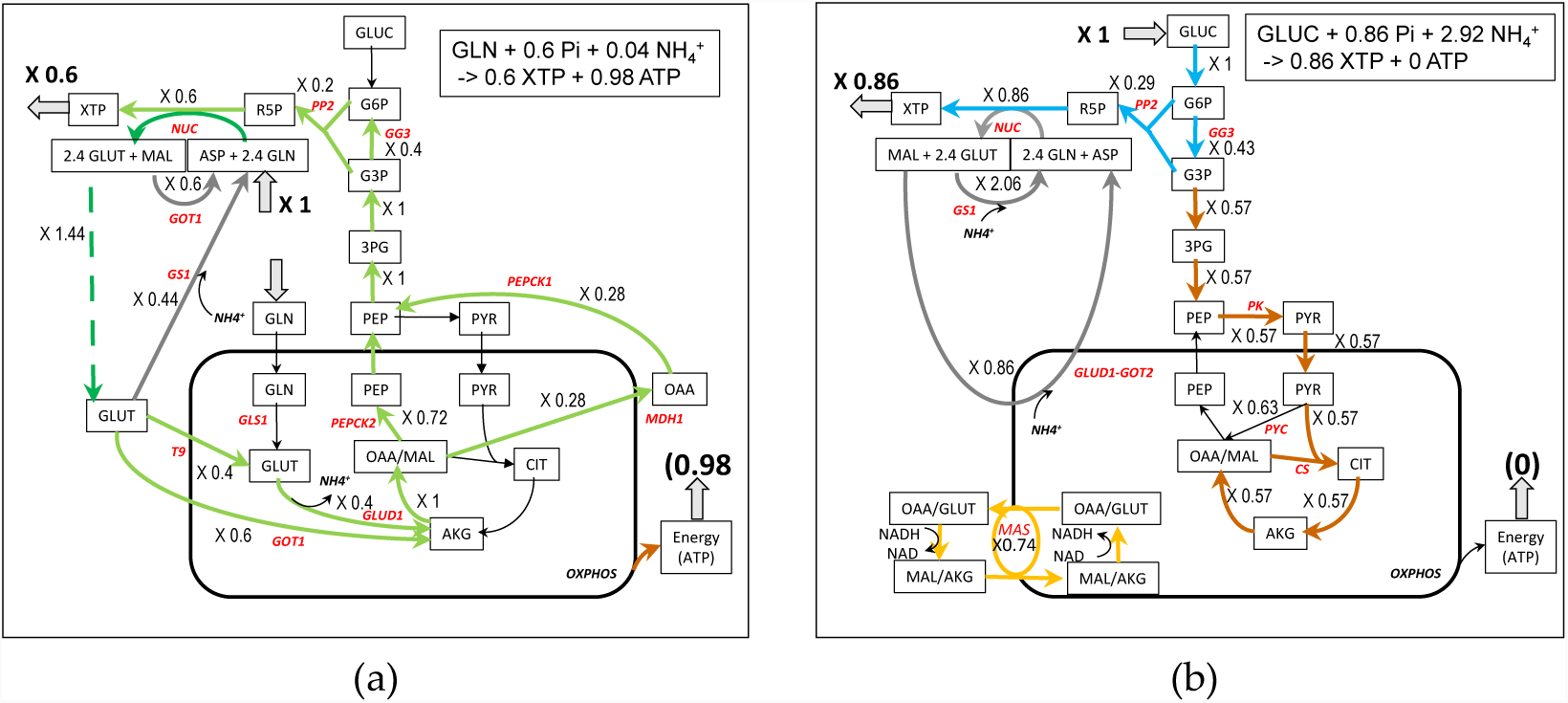
Simplified representation of nucleotides synthesis from glutamine (a) and glucose (b). Note that in (b), the flux through T4 and GOT 2 can be split, in the malate-aspartate shuttle (MAS) with a flux of 0.74 and in a 0.86 flux corresponding to aspartate recycling. Complete model in Figure S5 (a) and (b) in the supplementary materials.

The synthesis of nucleotides from glucose requires R5P synthesis and the recycling of glutamine and aspartate. R5P is produced by the pentose phosphate pathway. Glutamine is recycled from glutamate (+NH_4^+^_) by glutamine synthase (GS1) and aspartate is recycled from fumarate then malate and oxaloacetate. There is the need to reoxidise glycolytic NADHc and the NADHc produced by nucleotide synthesis inducing a 0.74 flux through the malate-aspartate shuttle (see Figures 5b and S5b). The yield of nucleotide synthesis with glucose is higher than with glutamine (0.86 XTP per glucose) but with a high input of NH_4^+^_ and no ATP production despite a higher activity of respiratory chain.

### 3.5 Fatty acids synthesis from glutamine and glucose (Figure 6)

Fatty acids (FA) synthesis is an important pathway in proliferating cells, especially for phospholipids syntheses. In the case of palmitate synthesis, 0.17 palmitate molecule can be synthesized from one molecule of glutamine and because this is rather energy consuming, no ATP molecule is synthesized (Figure 6a). The synthesis occurs through ATP citrate lyase (CL) fed by citrate synthesized in TCA cycle. The glutamine-derived AKG flux in the TCA cycle (flux = 1) is split between the reductive pathway giving directly citrate by inversion of IDH3 (flux = - 0.4) and the oxidative pathway through malate and oxalacetate (flux = 0.6) and pyruvate coming from the recycling of OAAc, product of CL in the cytosol. To these fluxes is superimposed a cycle involving IDH1 and IDH3 in order to transform NADHm into NADPHc necessary for the synthesis of fatty acids (flux = 2.4 in grey in figure 6a and S6a) giving a net reductive flux of IDH3 equal to – 2.8. NADPHc could have been made by the malic enzyme (ME1) in the cytosol but this occurs with a slightly lower yield (0.16 PAL per 1 glutamine). In [32] the authors showed that 60% of NADPH is synthesized from glutamine by malic enzyme and that the pyruvate produced is excreted as lactate. We do not observe a release of lactate in our models because, in the absence of glucose, there is no excess of carbon and pyruvate enters the mitochondria to replenish the TCA cycle in the classical anaplerotic way. The authors also observed a G6PDH flux (PP1) of the same order as the glutaminolysis flux, demonstrating that several sources of NADPH can be operating *in vivo* at the same time.

**Figure 6.**
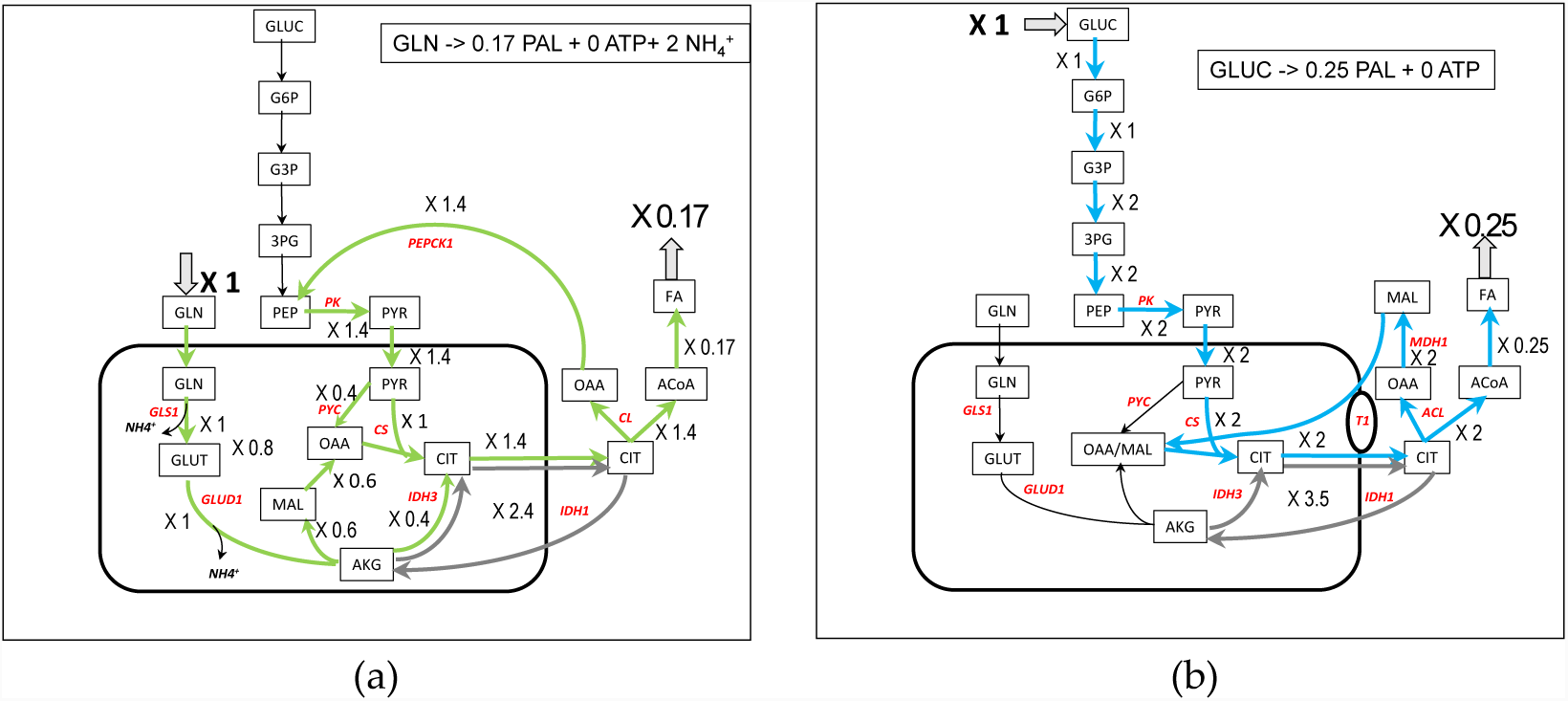
Simplified representation of palmitate synthesis from glutamine (a) and glucose (b). In grey, cycling of citrate accompanying the conversion of NADHm in NADPHc needed for palmitate synthesis. Complete model in Figure S6a and b in the supplementary materials.

The involvement of reductive use of glutamine and participation of IDH1 in fatty acids synthesis is well recognised [14,15,33] in hypoxia and with mutations in TCA cycle or in respiratory chain [34] and appears to be linked to the AKG/citrate ratio [35]. In our model (Figure 6a) the TCA cycle can be viewed as converging to citrate synthesis by the two oxidative and reductive pathways from AKG. This dual TCA pathway used for citrate synthesis is also well documented [36,37].

With glucose as carbon substrate (Figure 6b), oxalacetate coming from ATP-acetate lyase (CL) generates malate thanks to the cytosolic malate dehydrogenase (MDH1), oxidizing the glycolytic NADHc as found in [34,38]. The cytosolic malate re-enters mitochondria through antiporters, particularly in exchange with citrate. It is an elegant way to absorb the cytosolic reductive power of glucose avoiding the operation of MAS. This pathway is often observed when NADH reoxidation is impeded (respiratory complex deficiency for instance). Like with glutamine, there is a cycle involving IDH1 and IDH3 which transforms NADHm into NADPHc (flux = 3.5 in grey in figure 6b and S6b).

### 3.6 Serine synthesis from glutamine and glucose (Figure 7)

Serine is the major source of one-carbon units for methylation reactions via tetrahydrofolate and homocysteine. It is the precursor of glycine (see [39] for a mini-review). In his pioneer work, Snell demonstrated the importance of serine biosynthesis in rat carcinoma [40]. More recently several authors emphasized the role of serine in breast cancers and in melanoma cells [41–43].

**Figure 7.**
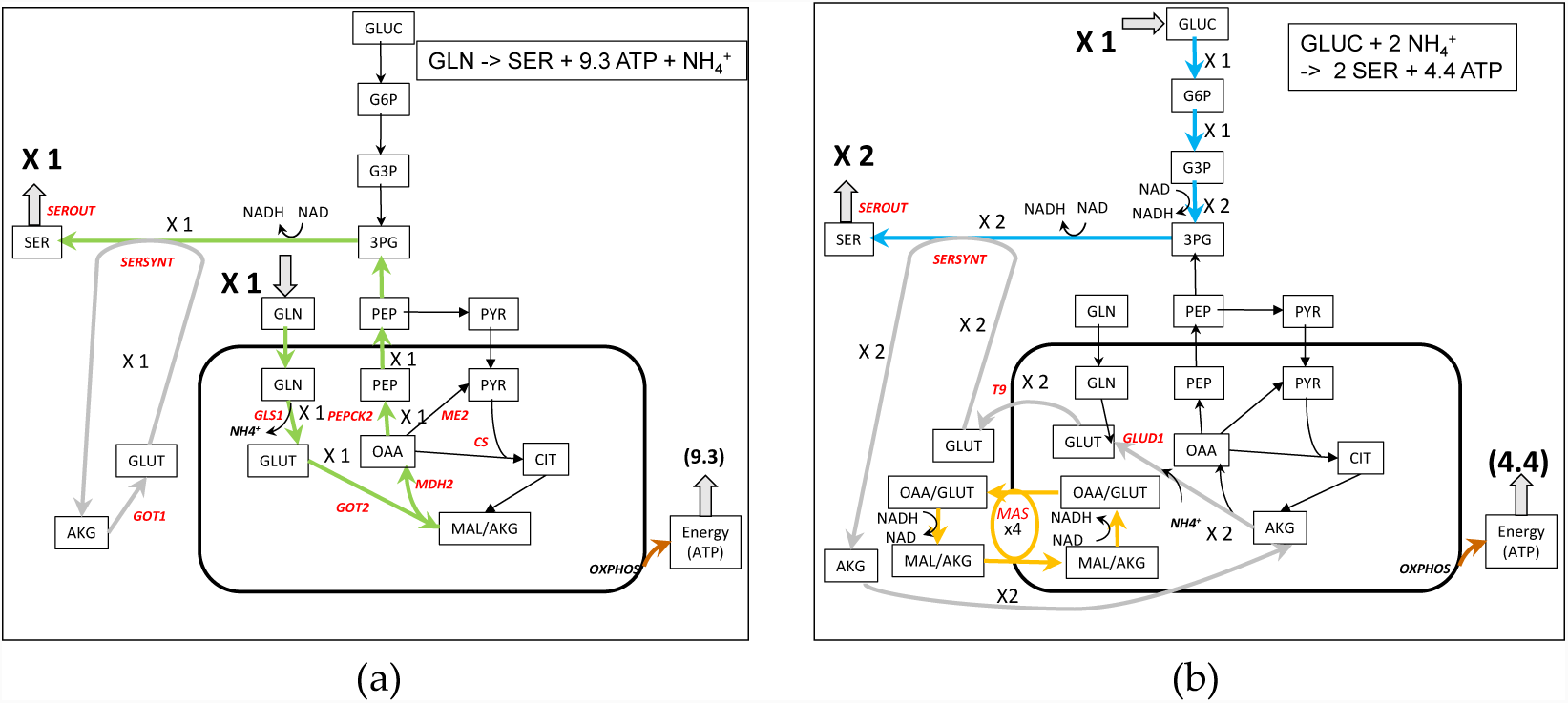
Simplified representation of serine synthesis from glutamine (a) and glucose (b). In light grey, the recycling of glutamate for serine synthesis. Complete model in Figures S7 in the supplementary materials.

We have already studied the different yield in serine synthesis on different substrates with our core model [44]. The synthesis of serine from glutamine is represented in Figure 7a. It involves the synthesis of PEP as in pyruvate synthesis, but here PEP is used to make 3PG, a serine precursor. The glutamate used in the transamination reaction is recycled from the AKG produced and the NADH produced by the 3-phosphoglycerate dehydrogenase reaction is reoxidized with MDH1. In addition, 9.3 ATP can be synthesized. With glucose as a carbon substrate, 3PG is synthesized directly from glucose via glycolysis (Figure 7b). However the work of Hanson in rats demonstrates that “pyruvate entry into the gluconeogeneic pathway is the major route for serine biosynthesis” [39,45]. It is difficult to obtain this pathway from glucose in C2M2N. This could indicate that in the case of this work, serine is mainly synthesized from other sources than glucose (glutamine or other amino acids for instance). The yield of serine synthesis with glucose as a carbon substrate can reach 2 molecules of serine per molecule of glucose with 4.4 ATP molecules produced. 4 NADHc are generated which requires a flux of 4 in the MAS (Figure 7b).

### 3.7 Comparison of C2M2N with MitoCore

Similar FBA simulations have been performed with MitoCore which is a more detailed model of metabolism. The results are nearly identical and are presented in Table 1. Note that the synthesis of nucleotides is not included in MitoCore so it is not possible to compare XTP synthesis in both models. The SBML file of the version of MitoCore used in this paper (with only input of glutamine or glucose) can be found in the supplementary materials

### 3.8 Distribution between glutamine and glucose for the biomass of proliferating cell. The nitrogen metabolism

Proliferating cells (cancer or immune cells) necessitate the input of NH_3_ group for the synthesis of amino acids, nucleic acids and polyamines. It is widely accepted that glutamine is the main donor of nitrogen for proliferating cell with a central role of glutamate. We have shown above that glutamine as well as glucose can allow the synthesis of most of the essential metabolites which constitute the biomass of proliferating cells and supply cell energy (ATP) [26]. In the tumour and immune cells microenvironment, glutamine and glucose are simultaneously present and the question arises of how they divide the task of feeding biomass. To answer this question we used the biomass as the specific objective function of proliferating cells. We let free the uptake of glucose and glutamine, only dictated by the optimization procedure of the biomass, but with an arbitrary limit of 1, to avoid unlimited amount of biomass. We obtained a metabolic pattern (figure 8a and S8a) in which the cell incorporated a large amount of glutamine not only for NH_4^+^_ feeding but also in part for carbon supply and around 80% of ATP to sustain biomass formation as noticed in [26] (See the ATPc and NADHm balance in figure S8a). This occurs with a large output of ammonia close to the input of glutamine which is larger than the need in NH_3_ for the biomass as already noted [14,32,46]. Although ammonia is toxic for the cell, it has been demonstrated that cancer cells accumulate ammonia in their microenvironment at rather high concentration [47,48]. In [47] the authors show that actually, cancer cells released ammonia when ammonia concentration in their microenvironment is low and uptaked ammonia for higher ammonia concentrations with a threshold around 1 mM. Figure 8a corresponds to low ammonia concentration in the microenvironment. An easy way to simulate the high concentration in the microenvironment associated with an ammonia input is to prevent ammonia output from the cells. In these conditions the uptake of glucose and glutamine to generate one unit of biomass are similar (0.762 and 0.883) (figure 8b and S8b). In this simulation 100% of ATP derives from glucose as calculated from the origin (glucose or glutamine) of NADHm (see Figure S8b). AKGm derives entirely from glutamate by transamination (GOT2). The main salient point of this solution is the recycling of NH_4^+^_ released by glutaminase and incorporated in part of the AKGm mainly by the reversion of glutamate dehydrogenase as shown experimentally by Spinelli et al. (see Fig. 1F in [47]) and also to a lower extent by carbamyl-phosphate synthesis (CPS1_OTC). No output of NH_4^+^_ is observed. Another advantage of NH_4^+^_ incorporation by glutamate dehydrogenase is the reoxidation of part of NADHm so that the glutamine per se does not contribute to the mitochondrial redox load (see the NADHm balance due to glucose or glutamine in Figure S8b).

**Figure 8.**
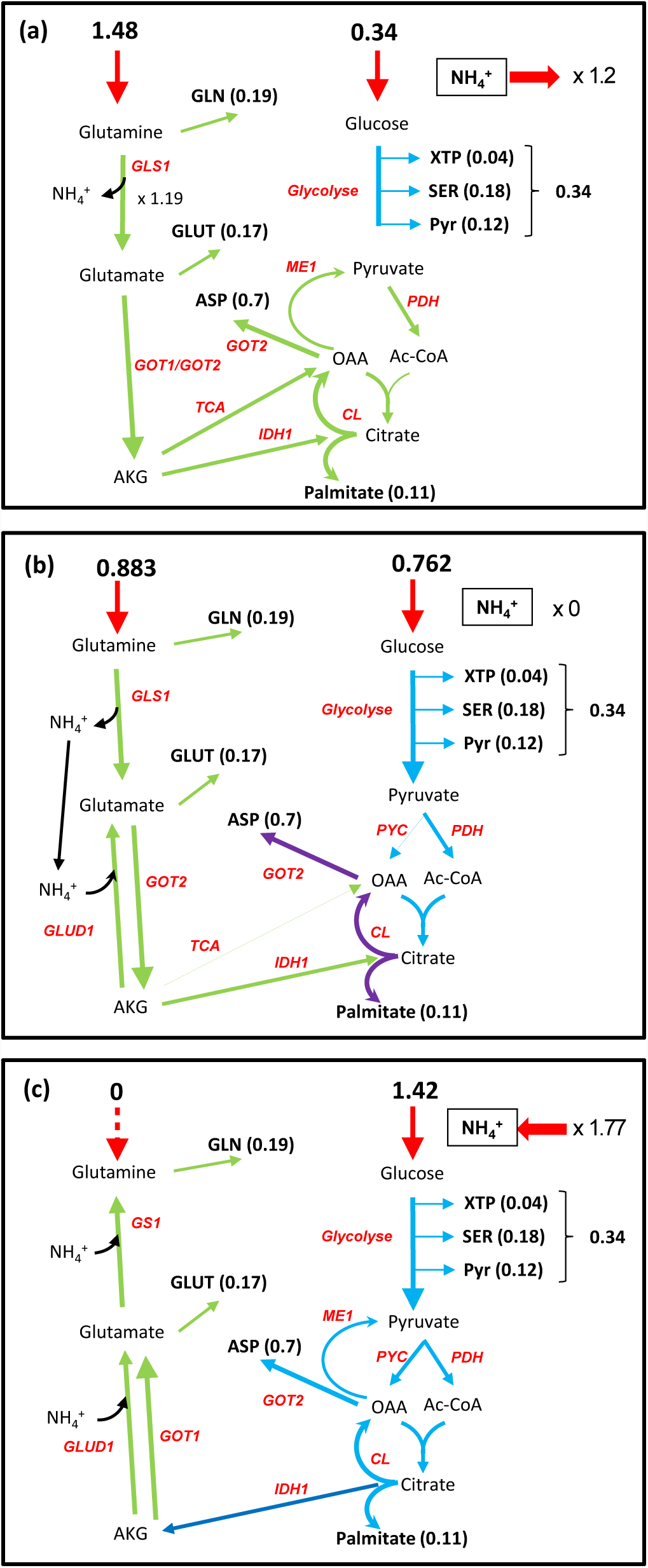
Interplay between glutamine, ammonia and glucose to sustain cell proliferation. (a) Nitrogen is incorporated as a large excess of glutamine with release of ammonia. (b) ammonia produced by GLS1 is stoichiometrically recycled by GLUD1(Glutamate dehydrogenase) in the reverse direction. In (c) Nitrogen is incorporated as ammonia in glutamate by the reversion of glutamate dehydrogenase and in glutamine by glutamine synthase. A small part of NH_4^+^_ is incorporated directly by CPS1 (carbamyl phosphate synthase). Complete metabolic networks in Figures S8a,b,c.

Glutamine limitation suppresses cancer cell proliferation which can be overcome by supplementing cells with ammonia [46,47]. It is easy to simulate this situation by suppressing the entry of glutamine inside the cells (Figure 8c and S8c). In these conditions the optimization of the biomass objective function relies on a massive entry of ammonia and glucose which is now the sole source of carbon and energy. As proposed in [49], this situation can occur when ammonia released locally by the catabolism of glutamine, diffuses in the microenvironment to cells for which the glutamine concentration in the microenvironment is lower.

If, using FVA, we look at essential reactions (those which are strictly positive or strictly negative and whose inhibition is susceptible to inhibit biomass formation) we obtain the reactions allowing biomass synthesis, i.e. CL, NUC, PL1, SERSYNT, ASS1_ASL_FH and T12 which represent more reactions due to the fact that most of these reactions stand for several consecutive reactions. All of which are therapeutically targeted in cancer [50]. Curiously GLS1 and GLUD1, the first steps in glutaminolysis, do not pertain to this set of reactions, indicating that other solutions which do not use these reactions exist.

If we put GLS1 to zero we obtain a solution in which NH_4^+^_ is incorporated from the microenvironment instead of glutamine by the reversion of Glud1 (similarly to Figure 8c and S8c). If in addition to GLS1 inhibition we add the inhibition of Glud1 we still have the normal biomass production of 1 with all glutamine metabolised in glutamate by the nucleotide synthesis NUC with excretion of nucleotides. In these conditions it could be therapeutically pertinent to inhibit simultaneously nucleotide synthesis and glutamate dehydrogenase Glud1.

### 3.9 Rewiring urea cycle metabolism

Urea cycle is another way to get rid of ammonia and to insure particular syntheses. The complete urea cycle takes place in the liver. However some urea cycle enzymes are expressed in different tissues, particularly in proliferating cells, in order to provide cells with arginine, ornithine, citrulline and polyamines. Overexpression of arginase 1 has been described in different types of tumours [51,52]. As pointed out in [53], by deprivation, “arginine becomes an essential amino acid generating a vulnerability that is utilized to treat cancer using arginine-deprivation agents”.

We can show (Figure 9) that increasing arginase activity while limiting the input of glutamine at the initial level induces a decrease in the biomass by diminishing the availability of arginine. However, glutamine can overcome arginine deficiency by a net arginine synthesis from glutamate conferring resistance to arginine deprivation with “higher glutamate dehydrogenase and glutaminase expression and preferential vulnerability to glutamine inhibitors” [54].

**Figure 9.**
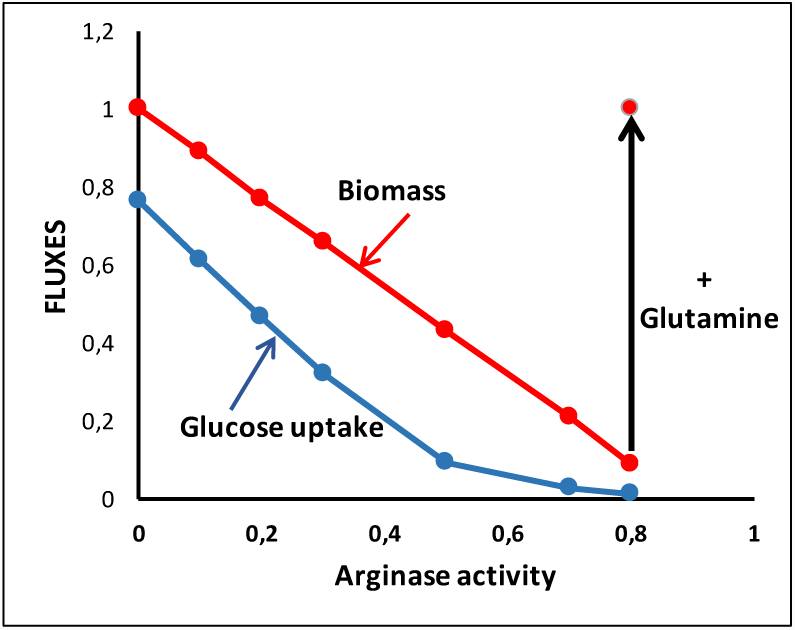
Arginine deprivation. Arginine deprivation is obtained with increased expression of Arginase in the situation of Figure 8b with the same input of glutamine (0.883). An increase in glutamine input (1.683) restore the biomass to 1.

### 3.10 Quantitative aspects of the Warburg effect

The Warburg effect or aerobic glycolysis is a salient feature of most cancer cells. It is characterized by an increase in glucose uptake and lactate release in the presence of oxygen. To simulate an increase in cell proliferation, we start from the situation of Figure 8a, i.e. a maximum of biomass equal to 1 with free availability of glucose and glutamine and we suppose that this situation corresponds to the maximum capacity of the respiratory chain (ETC flux = 2.374). Then we simulate an increase of biomass. The solution obtained is represented in Figure 10a. When biomass increases from 1, a massive glucose input is observed that supplements the extra need for ATP. It is accompanied by a massive release of lactate to reoxidize glycolytic NADH, a task that can no longer be provided by the respiratory chain. We also observe an increase of glutamine uptake (accompanied by an increase in NH_4^+^_ release) necessary for the biomass increase because the additional glucose carbons are lost with lactate excretion.

**Figure 10.**
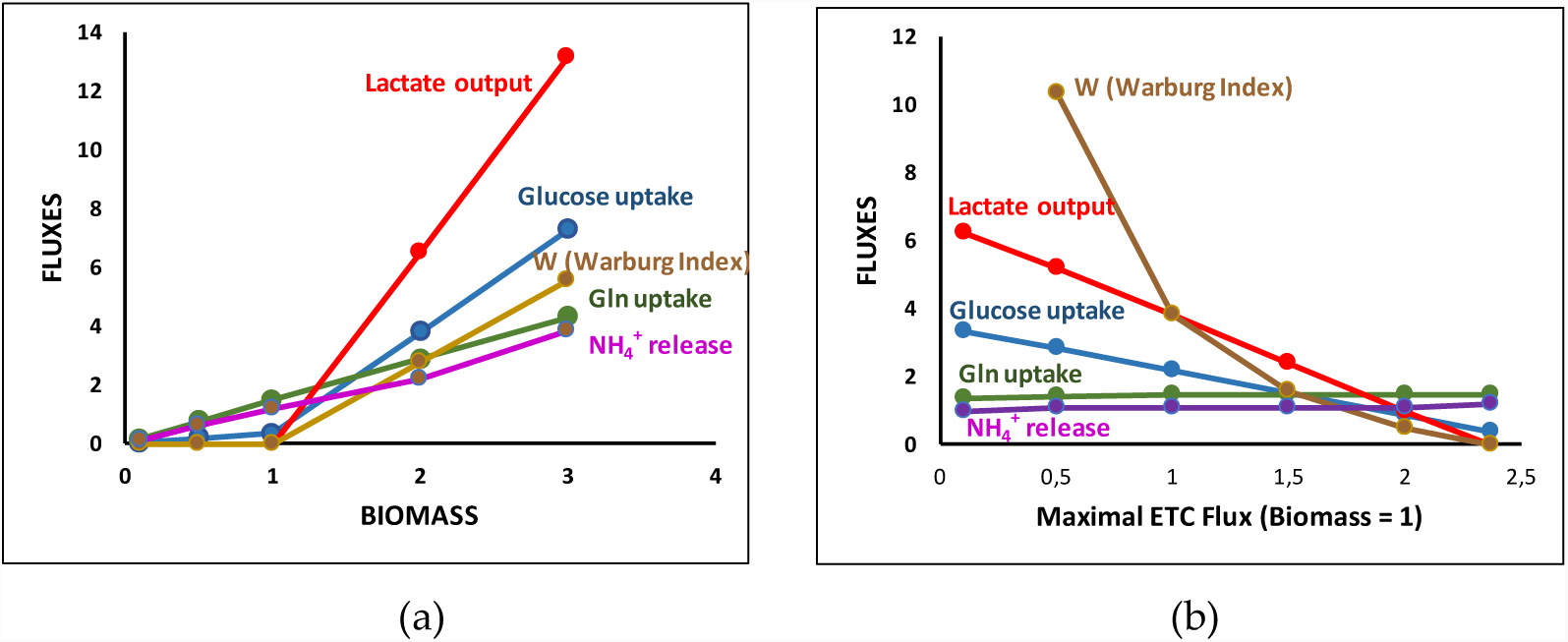
The Warburg effect. (a) Increase in biomass at constant respiratory chain content (ensuring maximum biomass equal to 1). (b) Decrease in respiratory rate from the sufficient value (2.374) to insure ATP synthesis and NADHc reoxidation. W is an index measuring the Warburg effect defined in [55] as the ratio of lactate release over respiratory chain (RC34) activity.

We can take this problem the opposite way (Figure 10b), still asking for a biomass equal to one but inhibiting the respiratory rate from its normal value, 2.374, which simulates hypoxia. As in Figure 10a, an increase in glucose uptake was observed that compensates the decrease in respiratory rate accompanied by a greater increase in lactate release but a slight decrease in glutamine.

As noticed in [29,55], the Warburg effect appears as a respiratory chain inadequacy (not necessary a decrease) in proliferating conditions leading to an increase of glycolytic flux to provide the requisite extra amount of ATP and consequently an increase in lactate output to reoxidize the extra amount of NADHc. Dai et al. [55] also emphasize the “redox balancing in leading to the Warburg effect and its correlation to proliferation rate”.

## 4. Conclusion

We developed a core metabolic model of central carbon and nitrogen metabolism, C2M2N, with a limited number of reactions (77) and of metabolites (54 internal metabolites) to explore the potential of glutamine to supplant glucose in metabolic syntheses, energy and biomass production. A salient feature of this model is that it takes into account the actual stoichiometries of the reactions particularly the stoichiometries of cofactors, even for concatenated reactions. A second characteristic of this model is to consider mitochondria as an isolated compartment with the relevant transporters. This led us to the third characteristics, to develop a relevant model of oxidative phosphorylation taking into account the mitochondrial ΔµH^+^ in the form of pseudo substrates DPH and DPSI as was already done in MitoCore [11] and in [2]. The last characteristic of C2M2N is that it takes into account nitrogen metabolism in addition to carbon metabolism. This led us to an easily tractable model that can produce rigorous quantitative simulations and testable predictions taking into account the constraints due to the regeneration of cofactors and a compromise between metabolite synthesis and energy production. Note that the model is not fixed; reactions can be added or cancelled to answer specific questions (for instance: introduction of DH reaction to simulate dehydrogenase activities able to regenerate NADc [29] in the synthesis of aspartate (Figures 4c and d)).

The advantage of C2M2N is that, due to the low number of reactions and metabolites, the interpretation of the results are rather straightforward and can be easily represented and understood on a metabolic scheme. All the solutions maximizing an objective function (described by FVA) can be more easily explored as done in supplementary material in the case of pyruvate synthesis and in section 3.8 in the case of biomass formation. Furthermore quantitative balance of any internal metabolites can be performed as in section 3.8 for biomass production. Furthermore, when glutamine and glucose are used together it is possible to distinguish which one contributes to any particular synthesis (see balance of NADHm due to glucose or glutamine in Figure S8 a and b for instance). Such a study makes it possible to identify the metabolic pathways at work for the synthesis of a given metabolite. Whenever possible, they are represented in our figures with different colours (green for glutamine and blue for glucose).

With C2M2N, we demonstrated that glutamine is a precursor as good as glucose for the syntheses of the main metabolites necessary for cell proliferation and energy production and we were able to give the quantitative yield in these productions (table 1). The same yields were obtained with MitoCore that constitutes a cross validation of both models. Taking into account cofactors made it possible to emphasize the role of the malate/aspartate shuttle (MAS) in glucose metabolism by making it a controlling (limiting) step in the use of glucose for metabolic syntheses and energy production, reorienting, when necessary, glycolysis towards lactate production (Crabtree or/and Warburg effect). This was also well exemplified by the “gas pedal” mechanism where Ca^2+^ activates the glutamate/aspartate carrier (T4) enhancing pyruvate formation and mitochondrial respiration [56]. This effect does not occur with glutamine, which can feed TCA cycle without NADHc production. More generally our study emphasizes the role of mitochondrial transporters which are often antiporters leading to metabolites cycling (coloured in grey in the figures) to output other metabolites. One can mention for instance glutamate cycling to remove mitochondrial aspartate or malate cycling to remove mitochondrial citrate (see Figure 6a and b and S6a and b for instance) and also the well-known malate-aspartate shuttle. These cycles could be controlling steps in metabolic network and thus good targets for therapeutic drugs.

C2M2N takes into account the constraints introduced by nitrogen metabolism which appear in the simultaneous presence of glucose and glutamine for biomass synthesis (section 3.8). We quantitatively evidenced the role of ammonia in the microenvironment and its possible recycling inside the same cell or between cells of the same tumour as recently demonstrated [47]. We also show the role of glutamine in complementing arginine deprivation as evidenced in [54] for instance.

In line with [55], we used C2M2N as a quantitative model to study the impact of a given amount of respiratory chain complexes on the relative uptake of glucose and glutamine and the release of lactate known as the Warburg effect. The Warburg effect clearly appears in our model as a way to resolve the tensions arising from an increased demand in ATP and the necessary reoxidation of the reducing power of proliferating cells. C2M2N allowed us to depict the causal link leading to the Warburg effect: when the activity of respiratory chain cannot be increased upon an increase in biomass production rate, the oxidative phosphorylation limits both the NADHm reoxidation and the percentage of mitochondrial ATP production. The only way to supply the extra demand in ATP is through glycolysis with an increased uptake of glucose but also an increased production of NADHc, which cannot be reoxidized by the respiratory chain. NADHc is thus reoxidized thanks to LDH with the release of lactate that dissipates the extra glucose carbons. Others sources of carbons, among which glutamine uptake, are thus necessary to sustain an increase in biomass production rate. In parallel, glutamine acts as an NH3 group donor but can also modulate the cell redox state through a balance between its oxidative or reducing catabolism. Of importance is the possible trend of glutamine to answer the increase in reducing power in making NADPH mainly through IDH1 or ME1; NADPH can be then reoxidized in fatty acids and glutathion syntheses.

C2M2N allowed us to look for the steps whose inhibition could decrease biomass production and thus cell proliferation. In our model, the inhibition of any pathway leading to a biomass component will inhibit cell proliferation. This is a good indication of the main targets for anticancer drugs which has nevertheless to be considered with caution because several other minor pathways may exist which are not considered in C2M2N but can be overexpressed in pathological cells. However, it appears in our model that the inhibition of the first steps of glutaminolysis, GLS1 and Glud1 (see section 3.8) are not sufficient to inhibit biomass formation. Several steps inhibitions are necessary to this aim. This point would be more evident on genome-scale models showing that C2M2N utilization does not exempt from the consideration of larger models. Nevertheless, C2M2N can constitute a first step to analyse experimental results with the quantitative constraints restricting the study to the possible relevant solutions. It can also be a first step in analysing the solutions of genome scale metabolic models by the recognition of common solutions and the reasons of non-common ones. Passing through a middle size metabolic model such as MitoCore can be an additional option with an increase by an order of magnitude in the number of reactions (77 reactions for C2M2N, 407 for MitoCore and around 7 440 for Recon 2 [4]).

A drawback of genome scale models and of large models in general (more than several hundreds of reactions) is that flux balance analysis is the only possible approach. The number of elementary flux modes (EFMs) is usually out of calculation, and taking into account the rate equations in differential equations reflecting the change in the concentration of metabolites faces a number of difficulties (their number, the knowledge of all kinetic constants, etc). Similarly metabolic control analysis (MCA) which can give an idea of pertinent therapeutic targets is unrealizable. These tasks are possible and easy with smaller metabolic models restricted to well-known reactions. The interest of having different approaches to explore a metabolic network is to be able to choose the most appropriate method for each particular biological question and more generally to confront the solutions given by the different theoretical approaches.

We think that C2M2N is a good compromise between a comprehensive compilation of all the metabolic steps and a much lower number of the most important reactions of central carbon and nitrogen metabolism that can give a realistic and quantitative representation of metabolism in many physiological or pathological conditions.

## Supporting information

xml file for MitoCore With glucose supply

xml file for C2M2N with gutamine supply

xml file for C2M2N with glucose supply

xml file for MitoCore With glutamine supply

## Supplementary Materials

Supplementary Materials 1: Others solutions for pyruvate synthesis from glutamine

Supplementary Materials 2: Decomposition of the metabolic network of Figure 8c in Elementary Flux Modes. Figure S1: C2M2N metabolic network.

Figure S2: complete C2M2N metabolic network of energy production from glutamine and glucose.

Figure S3: complete C2M2N metabolic network of pyruvate synthesis from glutamine and glucose.

Figure S4: complete C2M2N metabolic network of aspartate synthesis from glutamine and glucose.

Figure S5: complete C2M2N metabolic network of nucleotides synthesis from glutamine and glucose.

Figure S6: complete C2M2N metabolic network of Fatty Acids synthesis from glutamine and glucose.

Figure S7: complete C2M2N metabolic network of serine synthesis from glutamine and glucose.

Figure S8: complete C2M2N metabolic network of biomass production (BM = 1) with various amount of glutamine and glucose together.

Figure S9: complete C2M2N metabolic network of high biomass production (BM = 3) with various amount of glutamine and glucose together.

## Author Contributions

JPM and SR equally contributed to this work.

## Acknowledgments

We would like to thank several colleagues for helpful discussions on our model: Bertrand Beauvoit, Sophie Colombié, Anne Devin, Martine Dieuaide-Noubhani, Yves Gibon, Christine Nazaret, Pierre Petriacq, Sylvain Prigent and Michel Rigoulet. Thanks to Anne Devin and Louise Injarabian for English corrections.

## Appendix A: Abbreviations

ACS: Acetyl-CoA Synthase.
AGC: aspartate-glutamate carrier (see **T4** in appendix B)
AKG: α-ketoglutarate or 2-oxoglutarate
ANT: ADP/ATP exchanger.
ARGASE: arginase
ASS1_ASL_FH: argininosuccinate synthase + argininosuccinate lyase + fumarate hydratase
ASYNT: ATP Synthase.
ASPUP: Uptake of aspartate.
ATPASE: ATP usage.
BM: 0.18 SERc + 0.11 Palmitate_c + 0.04 XTPc + 0.19 GLNc + 0.17 GLUTc + 0.7 ASPc + 0.05 ARGc + 0.12 PYRc + 4 ATPc ==> 1 Biomass + 4 ADPc + 4 Pic.
CL (ACL): (ATP) Citrate Lyase.
CPS1_OTC: carbamoyl phosphate synthase 1 + Ornithine transcarbamylase
CS: Citrate Synthase.
DPH: pH difference between inside and outside mitochondria.
DPSI: mitochondrial membrane potential.
ENOMUT: Enolase + Phosphoglycerate Mutase.
ETC: Electron Transport Chain or Respiratory Chain
FBA: Flux Balance Analysis.
FVA: Flux Variability Analysis.
G1: hexokinase + phosphoglucose isomerase.
G2: phosphofructokinase + aldolase + triose-phosphate isomerase.
G3: Glyceraldehyde-3P Dehydrogenase + phosphoglycerate kinase.
GG3: triose phosphate isomerase + aldolase + fructose -1,6-biphosphatase.
GG4: phosphogluco isomerase + glucose-6-phosphatase.
GLS1: Glutaminase.
GLNUP: Uptake of Glutamine.
GLUCUP: Uptake of glucose.
GLUD1: Glutamate Dehydrogenase.
GOT: Glutamate Oxaloacetate Transaminase.
IDH: Aconitase + Isocitrate dehydrogenase.
L: Proton leak of the mitochondrial membrane.
LACIO: Input/Output of lactate.
LDH: Lactate Dehydrogenase.
MAS: Malate/Aspartate Shuttle.
ME: Malic Enzyme.
MDH: Malate Dehydrogenase
NAGS_ACY: N-acetylglutamate synthase + Amino Acylase (ornithine synthesis from glutamate).
NIG: for nigericine, exchange of DPH and DPSI.
NOS: NO synthase.
NUC: Nucleotide (XTP) Synthesis.
OGC: Oxoglutarate carrier (see **T2** in appendix B).
ORNT1: Ornithine/Citrulline + H+ exchanger.
ORNT2: Ornithine/ H+ exchanger.
PDH: Pyruvate Dehydrogenase.
PEPCK: PhosphoEnolPyruvate Carboxy Kinase.
PK: Pyruvate Kinase.
PL1: Synthesis of PhosphoLipids.
PP1: Oxidative part of PPP.
PP2: non-oxidative part of PPP.
PPP: Pentose Phosphate Pathway.
PYC: Pyruvate Carboxylase.
RC1: Complex I of Respiratory Chain.
RC2: succinate dehydrogenase + fumarase.
RC34: Complex III+IV of Respiratory Chain.
SEROUT: Output of serine.
SERSYNT: Serine Synthesis= Dehydrogenase + Transaminase and Phosphatase.
SLP (Substrate Level Phosphorylation): 2-oxoglutarate dehydrogenase + succinate thiokinase
XTP: Nucleotides

## Appendix B: METATOOL ENTRY FILE OF C2M2N

### -ENZREV

ANT ASS1_ASL_FH ASYNT ENOMUT G3 GLUD1 GOT1 GOT2 IDH1 IDH2 IDH3 LACIO LDH MDH1 MDH2 ME1 ME2 NIG NNT ORNT1 ORNT2 PP2 RC1 RC2 T1 T2 T3 T5 T7 T8 T9 T12 T14 T15 T16 T17 T20

### -ENZIRREV

ACS ARGASE ASPUP ATPASE BM CL CPS1_OTC CS G1 G2 GG3 GG4 GLNUP GLS1 GLUCUP GS1 L NAGS_ACY NOS NUC PDH PEPCK1 PEPCK2 PK PL1 PL2 PL3 PP1 PYC RC34 SEROUT SERSYNT SLP T4 T6 T11 T13 T18 T19

### -METINT

3PG ACETm ACoAc ACoAm ADPc ADPm AKGc AKGm ARGc ASPc ASPm ATPc ATPm CITc CITm CITRc CITRm DPH DPSI G3P G6P GLNc GLNm GLUCc GLUTc GLUTm LACc MALc MALm NADc NADHc NADm NADHm NADPc NADPm NADPHc NADPHm OAAc OAAm ORNc ORNm NH3 Palmitate_c PEPc PEPm Pic Pim PYRc PYRm R5P SERc SUCCm UREAc XTPc

### -METEXT

ARG ASP Biomass CO2 CoAc CoAm GLN GLUC GLUT HCO3 LAC NH3_out NO O PalCoAc PalCoAm Palmitate Pi PYR Q QH2 SER UREA XTP

### -CAT

ACS : ACETm + 2 ATPm + CoAm = ACoAm + 2 ADPm + 2 Pim.

ANT : ATPm + ADPc + DPSI = ATPc + ADPm.

ARGASE : ARGc = UREAc + ORNc. ASPUP : ASP = ASPc.

ASS1_ASL_FH : CITRc + ASPc + 2 ATPc = ARGc + MALc + 2 ADPc + 2 Pic.

ASYNT : 3 ADPm + 3 Pim + 8 DPH + 8 DPSI = 3 ATPm.

ATPASE : ATPc = ADPc + Pic.

BM : 0.18 SERc + 0.11 Palmitate_c + 0.04 XTPc + 0.19 GLNc + 0.17 GLUTc + 0.7 ASPc + 0.05 ARGc + 0.12 PYRc + 4 ATPc = 1 Biomass + 4 ADPc + 4 Pic.

CL : CITc + ATPc + CoAc = ACoAc + OAAc + ADPc + Pic.

CPS1_OTC : NH3 + HCO3 + ORNm + 2 ATPm = CITRm + 2 ADPm + 2 Pim. CS : ACoAm + OAAm = CITm.

ENOMUT : PEPc = 3PG.

G1 : GLUCc + ATPc = G6P + ADPc.

G2 : G6P + ATPc = 2 G3P + ADPc.

G3 : G3P + NADc + ADPc + Pic = 3PG + NADHc + ATPc.

GG3 : 2 G3P = G6P + Pic.

GG4 : G6P = GLUCc + Pic.

GLNUP : GLN = GLNc.

GLS1 : GLNm = GLUTm + NH3.

GLUCUP : GLUC = GLUCc.

GLUD1 : GLUTm + NADm = AKGm + NADHm + NH3.

GOT1 : GLUTc + OAAc = ASPc + AKGc.

GOT2 : GLUTm + OAAm = ASPm + AKGm.

GS1 : GLUTc + NH3 + ATPc = GLNc + ADPc + Pic.

IDH1 : CITc + NADPc = AKGc + NADPHc + CO2.

IDH2 : CITm + NADPm = AKGm + NADPHm + CO2.

IDH3 : CITm + NADm = AKGm + NADHm + CO2.

L : DPSI + DPH =.

LACIO : LACc = LAC.

LDH : PYRc + NADHc = LACc + NADc.

MDH1 : MALc + NADc = OAAc + NADHc.

MDH2 : MALm + NADm = OAAm + NADHm.

ME1 : MALc + NADPc = PYRc + NADPHc + CO2.

ME2 : MALm + NADm = PYRm + NADHm + CO2. NAGS_ACY : 2 GLUTm + ACoAm + NADHm + 3 ATPm = ORNm + CoAm + NADm + 3 ADPm + 3 Pim + AKGm + ACETm. NIG : DPSI = 4 DPH.

NNT : NADHm + NADPm + DPH + DPSI = NADm + NADPHm. NOS : ARGc + 1.5 NADPHc + 4 O = CITRc + 1.5 NADPc + NO.

NUC : R5P + 2 GLNc + ASPc + 6 ATPc + 0.2 NADc + 0.2 Q = XTPc + 2 GLUTc + 6 ADPc + 6 Pic + 0.2 NADHc + 0.2 QH2.

ORNT1 : ORNc + CITRm = ORNm + CITRc + DPH.

ORNT2 : ORNc = ORNm + DPH.

PDH : PYRm + NADm = ACoAm + NADHm + CO2. PEPCK1 : OAAc + ATPc = PEPc + ADPc + CO2.

PEPCK2 : OAAm + ATPm = PEPm + ADPm + CO2. PK : PEPc + ADPc = PYRc + ATPc.

PL1 : 8 ACoAc + 7 ATPc + 14 NADPHc + 7 HCO3 = Palmitate_c + 7 ADPc + 7 Pic + 14 NADPc + 8 CoAc + 7 CO2.

PL2 : PalCoAm + 7 NADm + 7 CoAm + 7 Q = 7 NADHm + 8 ACoAm + 7 QH2.

PL3 : Palmitate_c + CoAc = PalCoAc.

PP1 : G6P + 2 NADPc = R5P + 2 NADPHc + CO2.

PP2 : 3 R5P = 2 G6P + G3P.

PYC : PYRm + HCO3 + ATPm = OAAm + Pim + ADPm.

RC1 : NADHm + Q = NADm + QH2 + 4 DPH + 4 DPSI.

RC2 : SUCCm + Q = MALm + QH2.

RC34 : QH2 + O = Q + 6 DPH + 6 DPSI.

SEROUT : SERc = SER.

SERSYNT : 3PG + GLUTc + NADc = SERc + AKGc + NADHc + Pic.

SLP : AKGm + NADm + Pim + ADPm = SUCCm + NADHm + CO2 + ATPm.

T1 : CITm + MALc = CITc + MALm + DPH.

T2 : AKGc + MALm = AKGm + MALc.

T3 : MALm + Pic = MALc + Pim.

T4 : GLUTc + ASPm + DPH + DPSI = GLUTm + ASPc.

T5 : Pic + DPH = Pim.

T6 : PYRc + DPH = PYRm.

T7 : PEPm + CITc + DPH + DPSI = PEPc + CITm.

T8 : GLNc = GLNm.

T9 : GLUTc + DPH = GLUTm.

T11 : PalCoAc = PalCoAm.

T12 : Pi = Pic.

T13 : XTPc = XTP.

T14 : ASPc = ASP. T

15 : GLUTc = GLUT.

T16 : PYRc = PYR.

T17 : Palmitate_c = Palmitate.

T18 : ARGc = ARG.

T19 : UREAc = UREA.

T20 : NH3_out = NH3.

---

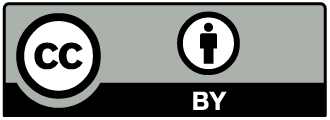

© 2018 by the authors. Submitted for possible open access publication under the terms and conditions of the Creative Commons Attribution (CC BY) license (http://creativecommons.org/licenses/by/4.0/).

## Supplementary Materials 1

Others solutions for pyruvate synthesis from glutamine. Multiplicity of solutions.

The solution obtained with C2M2N for the maximum rate of pyruvate synthesis from glutamine requires the maximum ATP synthesis flux. In the absence of this constraint, the emerging solution (with absolute fluxes minimization) is quite different (Figure S3c). Indeed, in this solution, pyruvate is synthesized through two pathways. The glutamine derived AKG engages in TCA cycle in both the reducing (flux = 0.61) and the oxidizing (flux = 0.39) pathways. The reducing pathway gives 0.61 citrate, which are split by ATP citrate lyase (CL) to give OAAc and acetyl-CoA ending to 0.077 palmitate. OAAc generates PEPc (PEPCK1) and then 0.61 pyruvate by PK. The 0.39 oxidizing pathway gives MALm which goes out to give MALc and then 0.39 pyruvate through ME1. There is also a 0.15 cycle involving MDH2 and ME1 which transforms 0.15 NADHm in 0.15 NADPHc to complete the 1.078 NADPHc (IDH1 and ME1) necessary for the synthesis of 0.077 palmitate. In this scenario, no ATP is synthesized and the reductive power of glutamine is consumed in citrate and palmitate synthesis. The mitochondrial production of NADH is low as shown by the low activity of RC1 (0.084) as compared with 3.0 in the previous solution allowing a high ATP synthesis flux.

Using MitoCore, we obtained a slightly different result with 1.41 pyruvate from glutamine and no ATP. In this solution, a great part of pyruvate is obtained with the successive syntheses of serine, glycine, alanine and then pyruvate. In this pathway, there is a high activity of monoamine oxidase specific of the central nervous system and of diabetic myocardium and a high activity of a mitochondrial lactate dehydrogenase which is rather unexpected in mammalian cells. Reducing mitochondrial lactate dehydrogenase (R_LDH_Dm_MitoCore) activity to zero gives a similar solution as the solution of C2M2N with no ATP synthesis above and a null activity of RC1 indicating a fine balance of oxidative and reductive glutaminolysis (IDH1 = −0.1 and IDH2 = − 0.495) in this case. A maximal ATP synthesis flux of 10.9 can be obtained when maintaining the maximal production of 1 pyruvate for 1 glutamine, which is comparable with the yields obtained with C2M2N. The activity of the respiratory chain is also the same as in C2M2N (R_CI = 3, R_CII = 1 and R_CIII = 4, MitoCore notations). However with MitoCore, pyruvate is made through three different pathways. Glutamine is first transformed in AKGm then in MALm (flux = 1). From there three fluxes will synthesize cytosolic pyruvate. The main one (flux = 0.8) goes through mitochondrial malic enzyme (ME2) giving PYRm released from mitochondria through an alanine cycle catalysed by the cytosolic and mitochondrial operation of ALATm and ALATc (alanine aminotransferase or glutamate-pyruvate aminotransferase, equivalent to GOT1 and 2 with ALA in place of ASP and PYR in place of OAA, not represented in Figure S3d). A second pathway (flux = 0.1) goes through MDH2, PEPCK2, PEP/Citrate exchanger and then pyruvate kinase. In the last pathway (flux = 0.1) MALm goes out with malate / phosphate exchanger and gives pyruvate through cytosolic malic enzyme (ME1).

These examples show the diversity of solutions, which can be explored through FVA (Flux Variability Analysis). Furthermore, with C2M2N, but not so easily with MitoCore, we can analyse each solution in terms of separate pathways and transporting cycles (in fact elementary flux modes (EFMs)). It is worth noting that we can represent the MitoCore solution on the C2M2N network with some adaptations.

## Supplementary Materials 2

Decomposition of the metabolic network of Figure 8c in Elementary Flux Modes.

The production of biomass is in blue with XTP, SER and PYR produced by the glycolysis. The other metabolites of the biomass (ASP, GLUT, GLN, ARG and PAL) are produced through citrate synthesis in mitochondria fed by pyruvate carboxylase. This is not so usual except for palmitate synthesis. The output flux of citrate is 1.7 which split in 0.88 for palmitate synthesis and 0.88 OAA precursor of the 0.7 ASP. The rest of citrate (0.8) is metabolized through IDH1 (giving a great part of the NADPH necessary for palmitate synthesis) to give AKG and GLUT (GOT1) and GLN (GS1). The rest of glutamate re-enter the mitochondria to make minute amount of ARG.

These fluxes are completed by several cycles necessary for the exchanges between mitochondria and cytosol. These cycles share some reactions so that it is not easy to decompose all the fluxes in a sum of cycles. Furthermore the decomposition is not necessarily unique. One decomposition is the following: the two cycles composing the MAS with a 2.7 flux; a AKGm-GLUTm-GLUTc cycle involving GOT2 (-Glud1) (-T9) T4 with a 1.33 flux. In this cycle, the aspartate and arginine NH3 are incorporated by (-Glud1). Then we have a 1.027 cycle GLUTc-GLUTm-AKGm-AKGc with T4 and T2. Then we have a 0.391 citrate cycle: CITm-CITc-AKGc-GLUTc-GLUTm-AKGm through (-IDH3) T1 IDH1 T4 GOT2. All these cycles superimpose to give the flux values (rounded values) indicated on Figure 8c.

**Figure S1.**
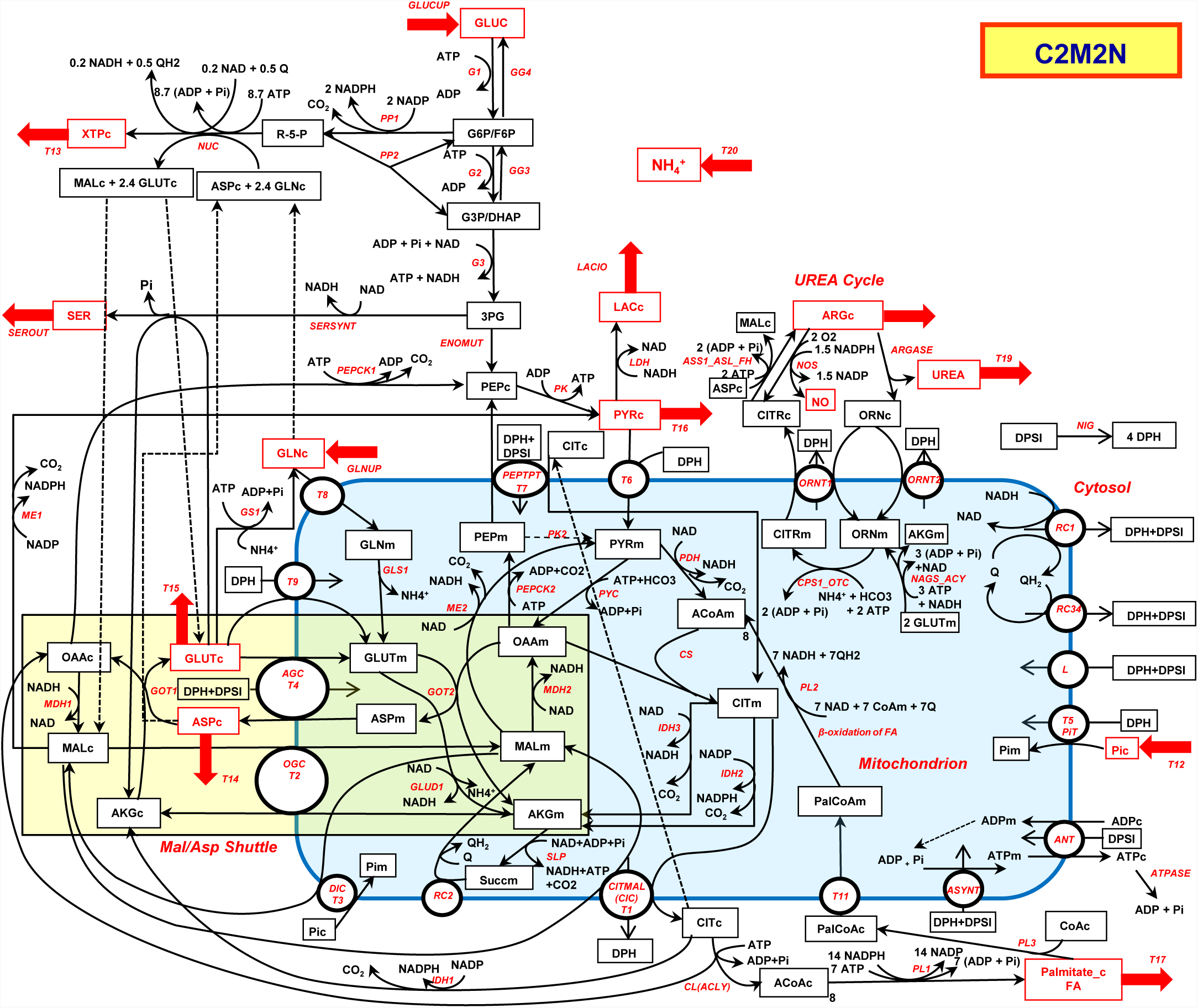
Identical to Figure 1 in the main text

**Figure S2a.**
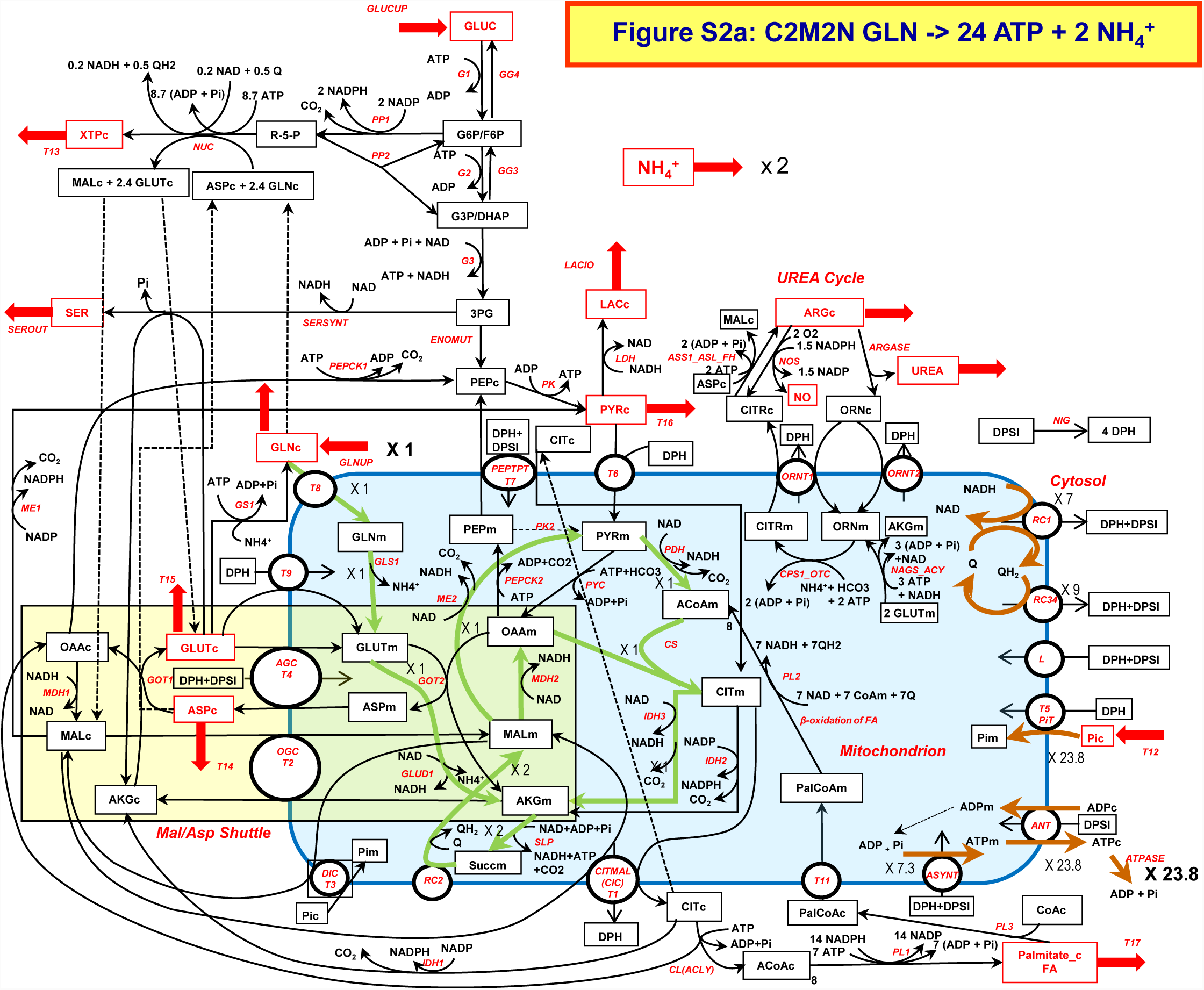
ATP production from glutamine. Green arrows represent the main pathway (flux = 1). Brown arrows depict OXPHOS. A complete TCA cycle is fed by α-ketoglutarate (AKG).

**Figure S2b.**
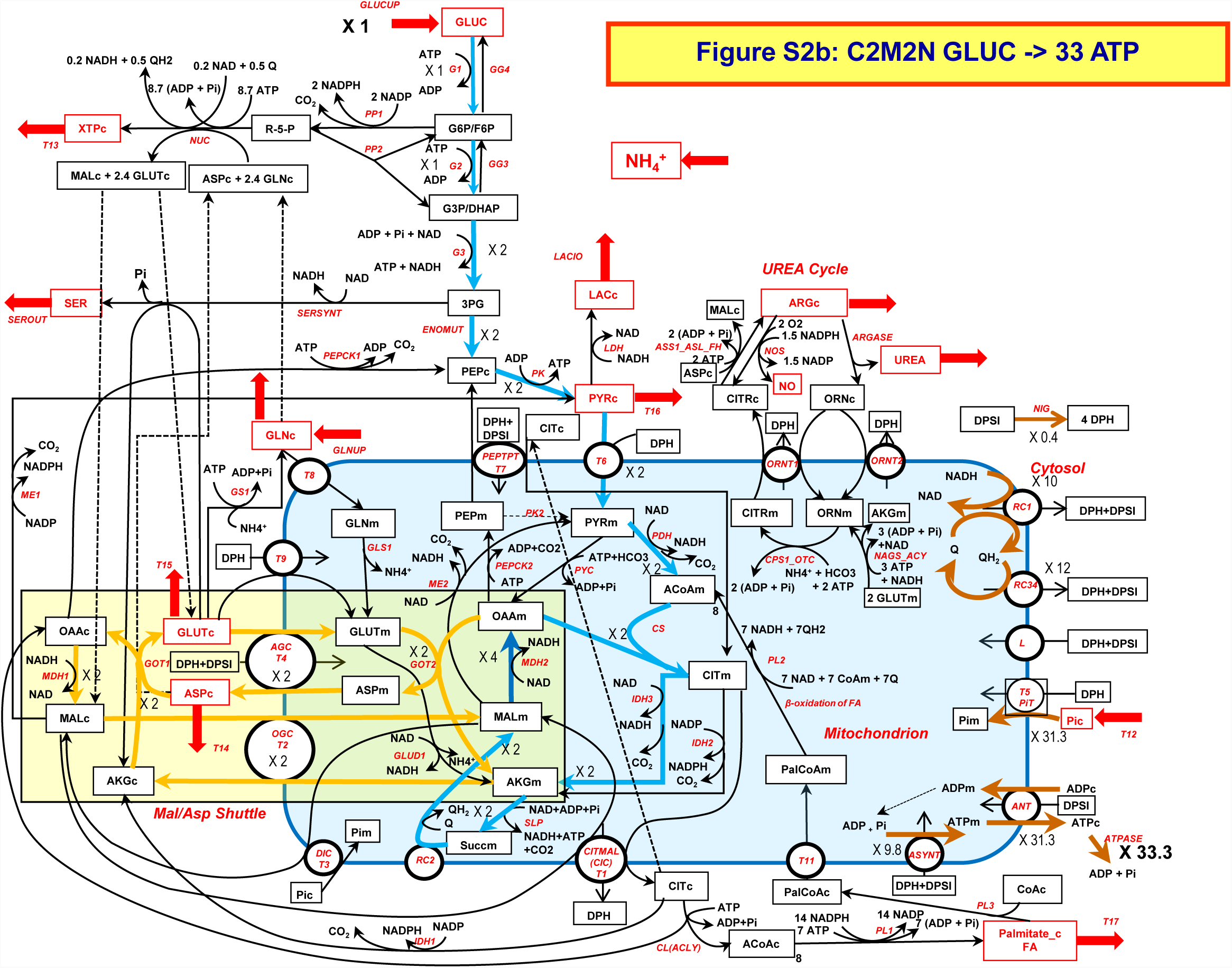
ATP production from glucose. Blue arrows represent the main pathway (flux = 2). Yellow arrows correspond to the malate/aspartate shuttle (flux = 2) to reoxidise glycolytic NADHc. Dark blue arrow means both blue and yellow pathway. Brown arrows depict OXPHOS

**Figure S3a.**
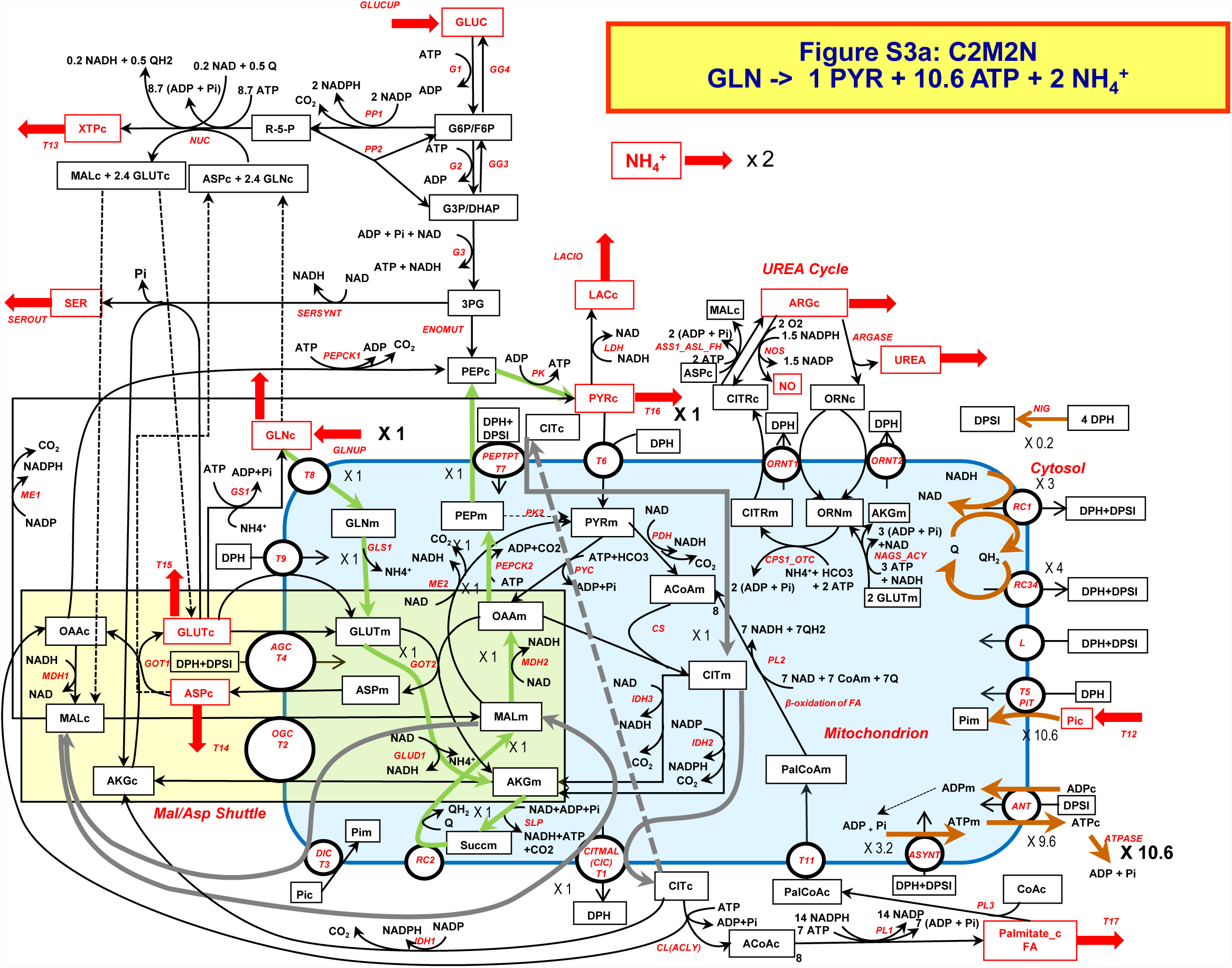
Pyruvate synthesis from glutamine with ATP production. Green arrows represent the main pathway (flux = 1). Brown arrows depict OXPHOS. Grey arrows depict a malate and citrate cycle to allow PEP output from mitochondria.

**Figure S3b.**
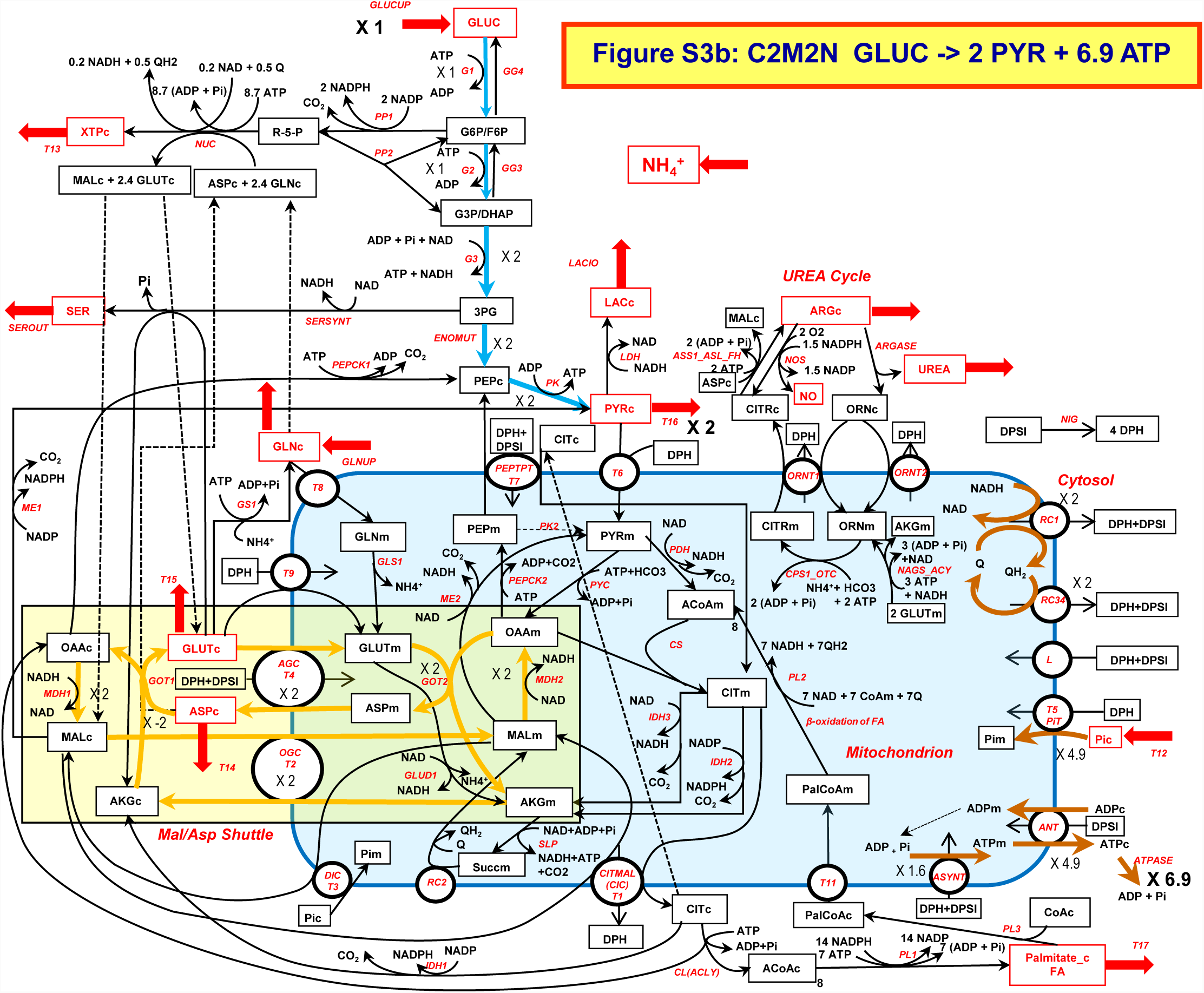
Pyruvate synthesis from glucose. Blue arrows represent the main pathway (flux = 2). Yellow arrows correspond to the malate/aspartate shuttle (flux = 2) to reoxidise glycolytic NADHc. Brown arrows depict OXPHOS

**Figure S3c.**
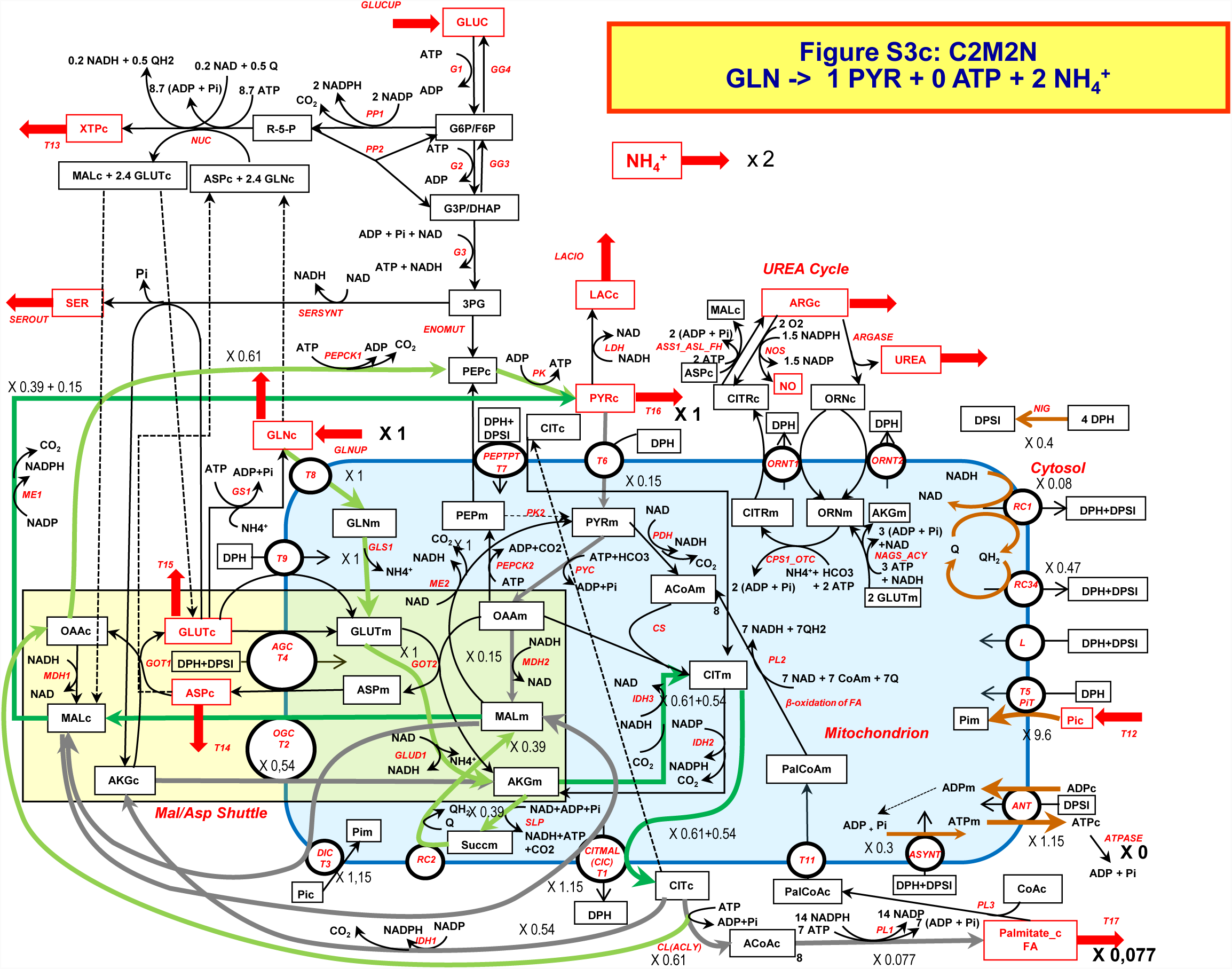
Pyruvate synthesis from glutamine without ATP production but with palmitate synthesis. Green arrows represent the main entry pathway (flux = 1) of glutamine GLS1 and GLUD1). The pyruvate synthesis is in green and the transporting cycles in grey. Pyruvate is synthesized by two pathways: a reducing pathway through IDH3 (dark green arrows= light green + grey fluxes) and an oxidizing pathway through MAL and ME1. Brown arrows depict OXPHOS. Note the low RC1 activity reflecting a low NADHm synthesis. The reducing power of glutamine is transformed in fatty acids (palmitate). See the supplementary Materials for a full description of this network.

**Figure S3d.**
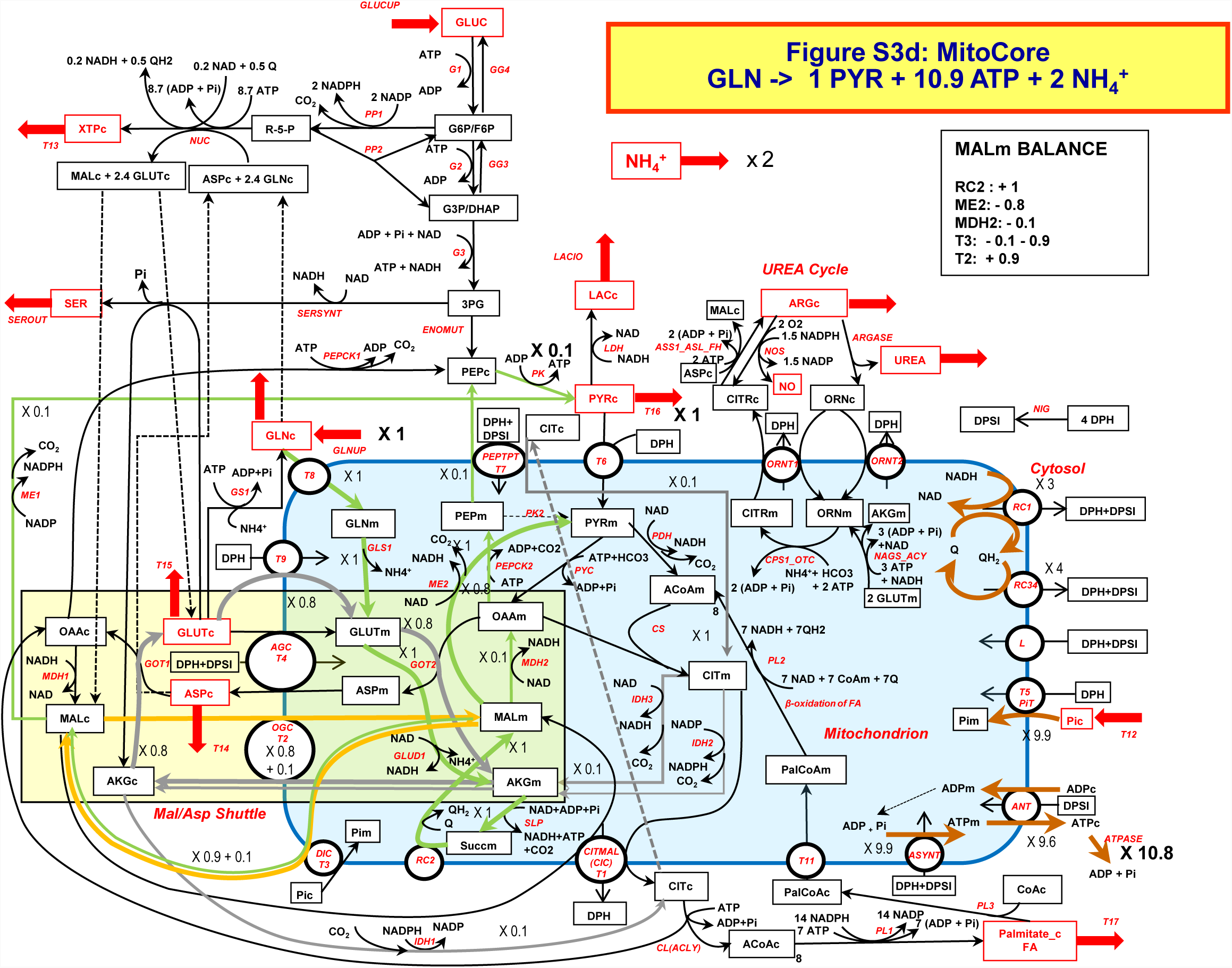
Pyruvate synthesis from glutamine with ATP production using MitoCore. Green arrows represent the three pathways of pyruvate synthesis from glutamine (total flux = 1). The main flux (thick green arrows) goes through ME2 and an alanine transporter not represented here. Two other fluxes (equal to 0.1 each; thin green arrows) go, one through MDH2, PEPCK2, T7 and PK; the other one goes through T3 and ME1. Brown arrows depict OXPHOS. Thin grey arrows (flux = 0.1) depicts a citrate/AKG cycle for release of PEPm in cytosol and thick grey arrows (flux = 0.8) a AKG/glutamate cycle contributing to alanine transport in cytosol. The yellow arrows depict a malate cycle for AKG transport in the grey AKG/glutamate cycle above.

**Figure S4a.**
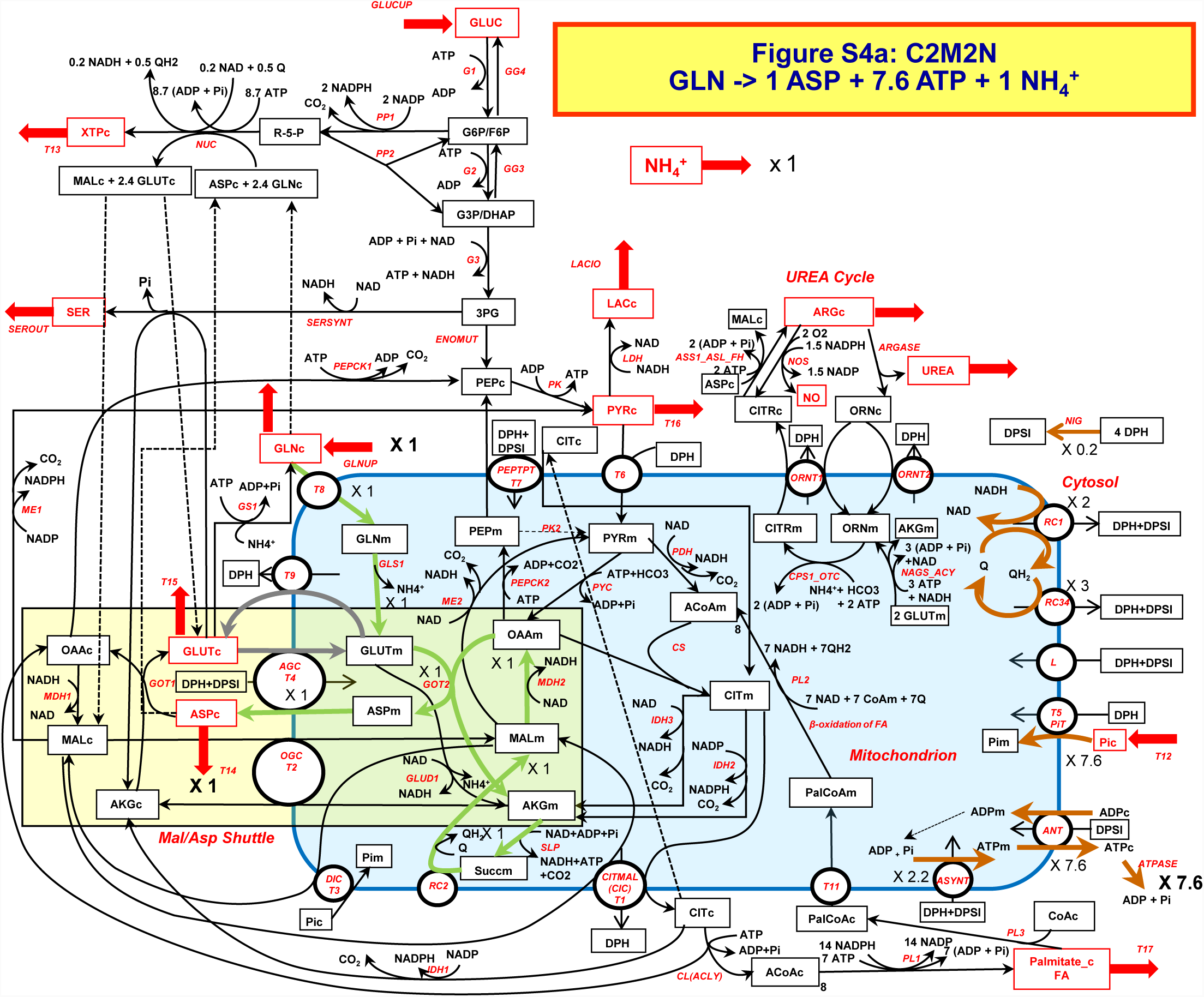
Aspartate synthesis from glutamine. Green arrows represent the main pathway (flux = 1). Grey arrows represent a glutamate cycle to output aspartate. Brown arrows depict OXPHOS

**Figure S4b.**
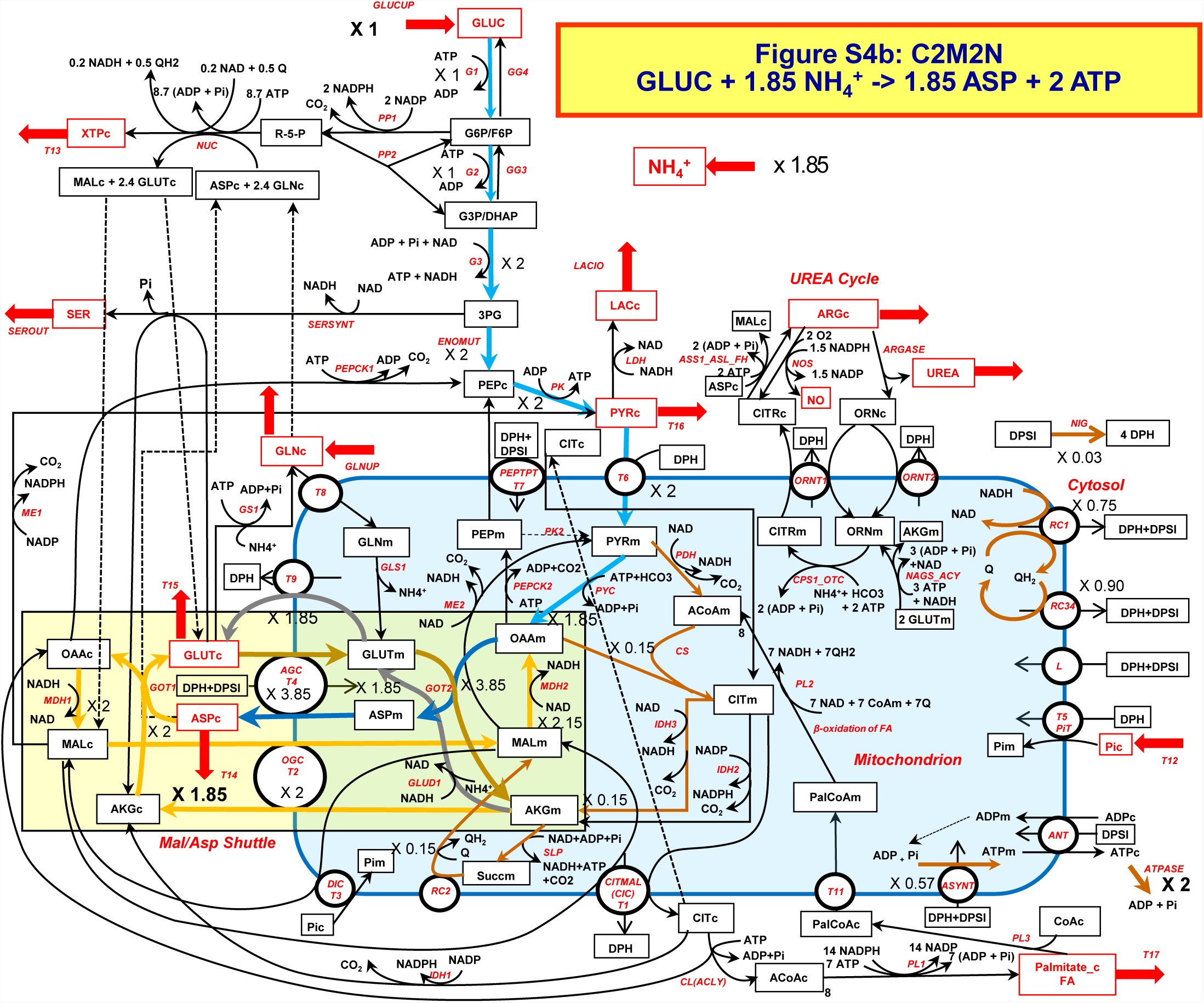
Aspartate synthesis from glucose with production of 2 ATP. Blue and grey arrows represent the aspartate synthesis and output (glutamate cycle) pathways (flux = 1.85). Yellow arrows correspond to the malate/aspartate shuttle (flux = 2) to reoxidise glycolytic NADHc. Dark blue and yellow arrows correspond to mixed pathways (MAS with blue or grey pathway). Pathway in brown arrows (TCA cycle = 0.15 and OXPHOS) are necessary for ATPm synthesis for PYC.

**Figure S5a.**
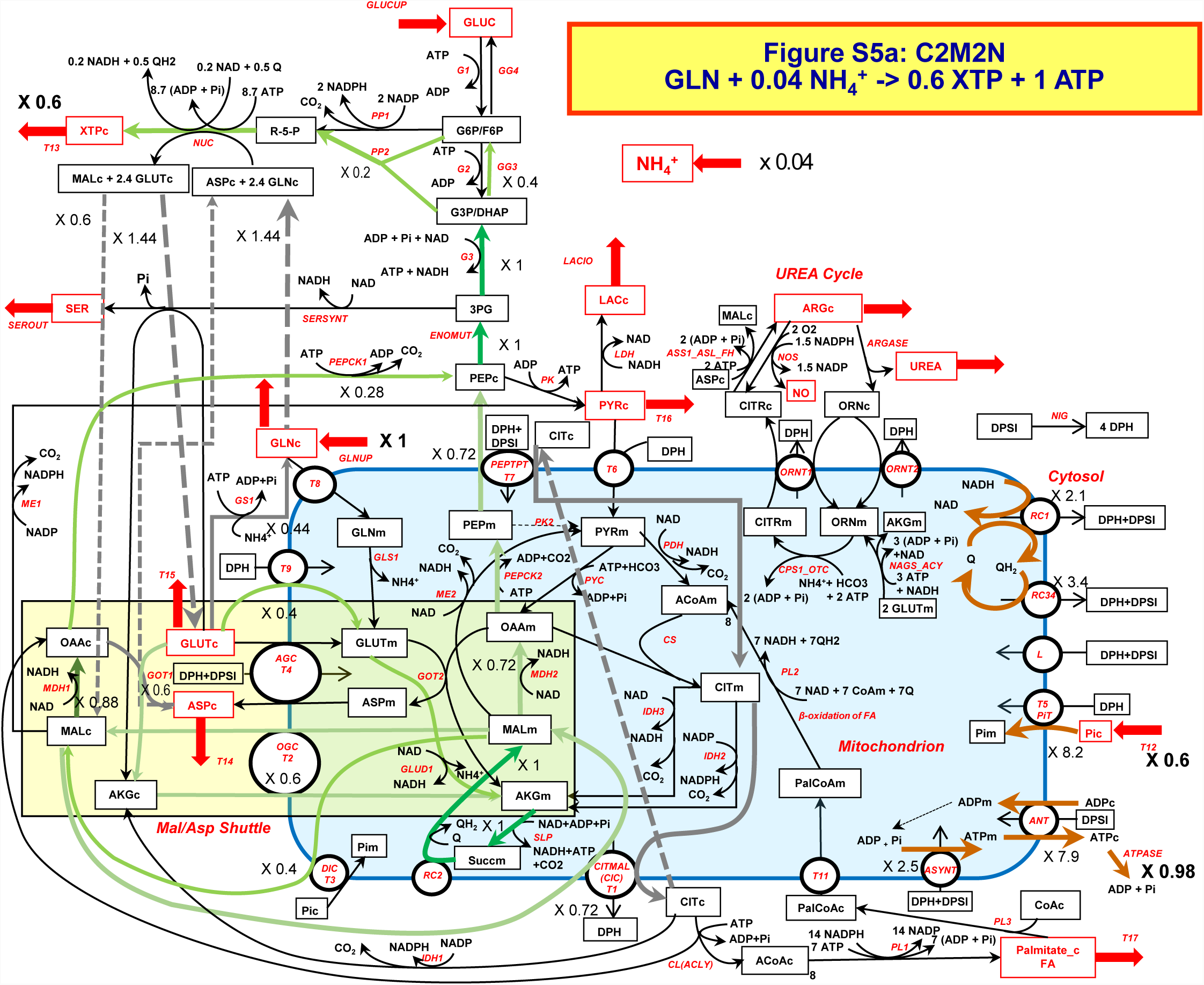
Nucleoside synthesis from glutamine. Green arrows represent the main pathway. Brown arrows depict OXPHOS. Grey arrows represents a citrate cycle necessary to export PEP from mitochondria.

**Figure S5b.**
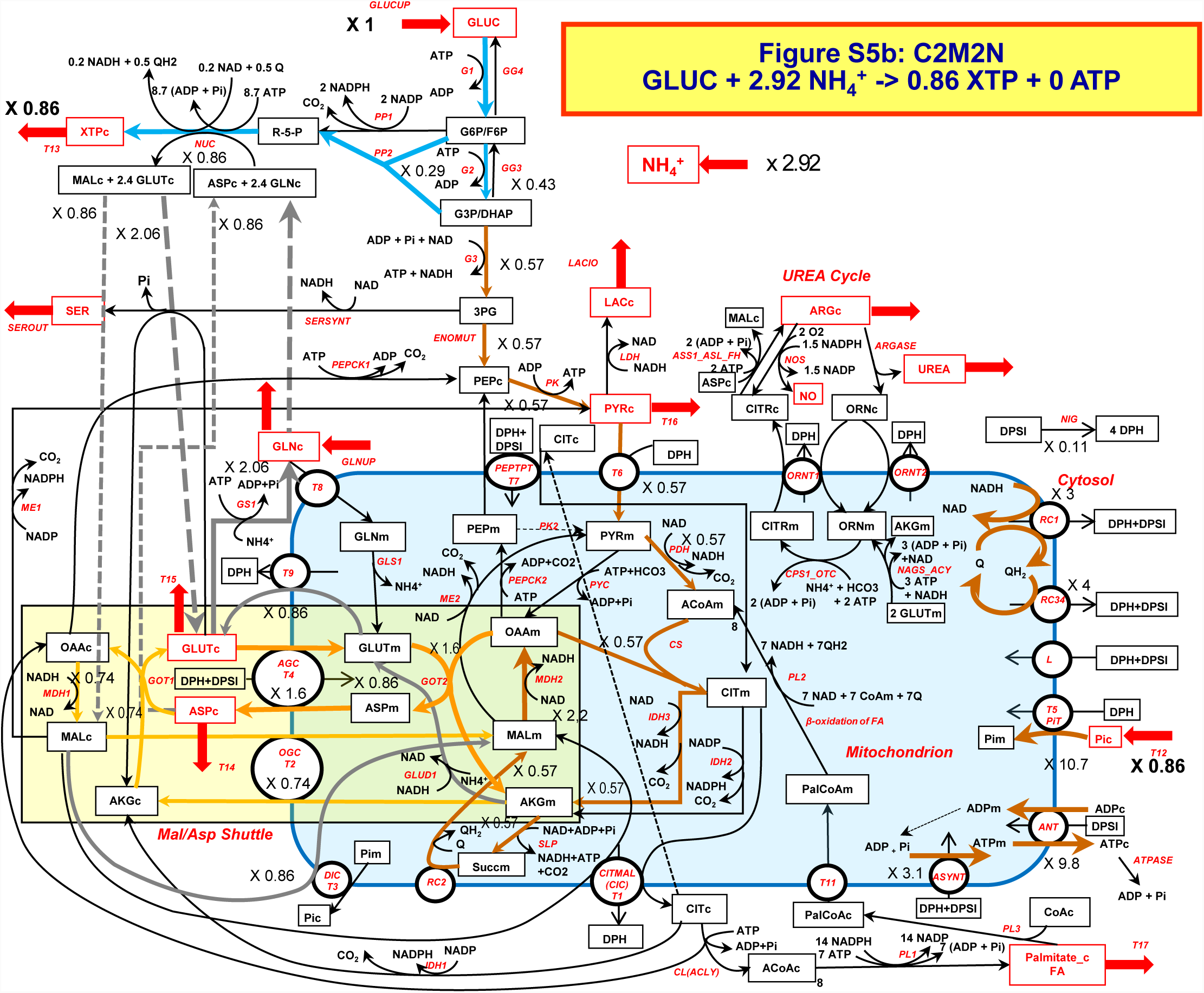
Nucleoside synthesis from glucose. Blue arrows represent the main pathway. Yellow arrows correspond to the malate/aspartate shuttle (flux = 2) to reoxidise glycolytic NADHc. Pathway in brown arrows (end of glycolysis + TCA cycle = 0.57 and OXPHOS) are entirely used to energy production (ATP necessary for (nucleotides synthesis (NUC)).

**Figure S6a.**
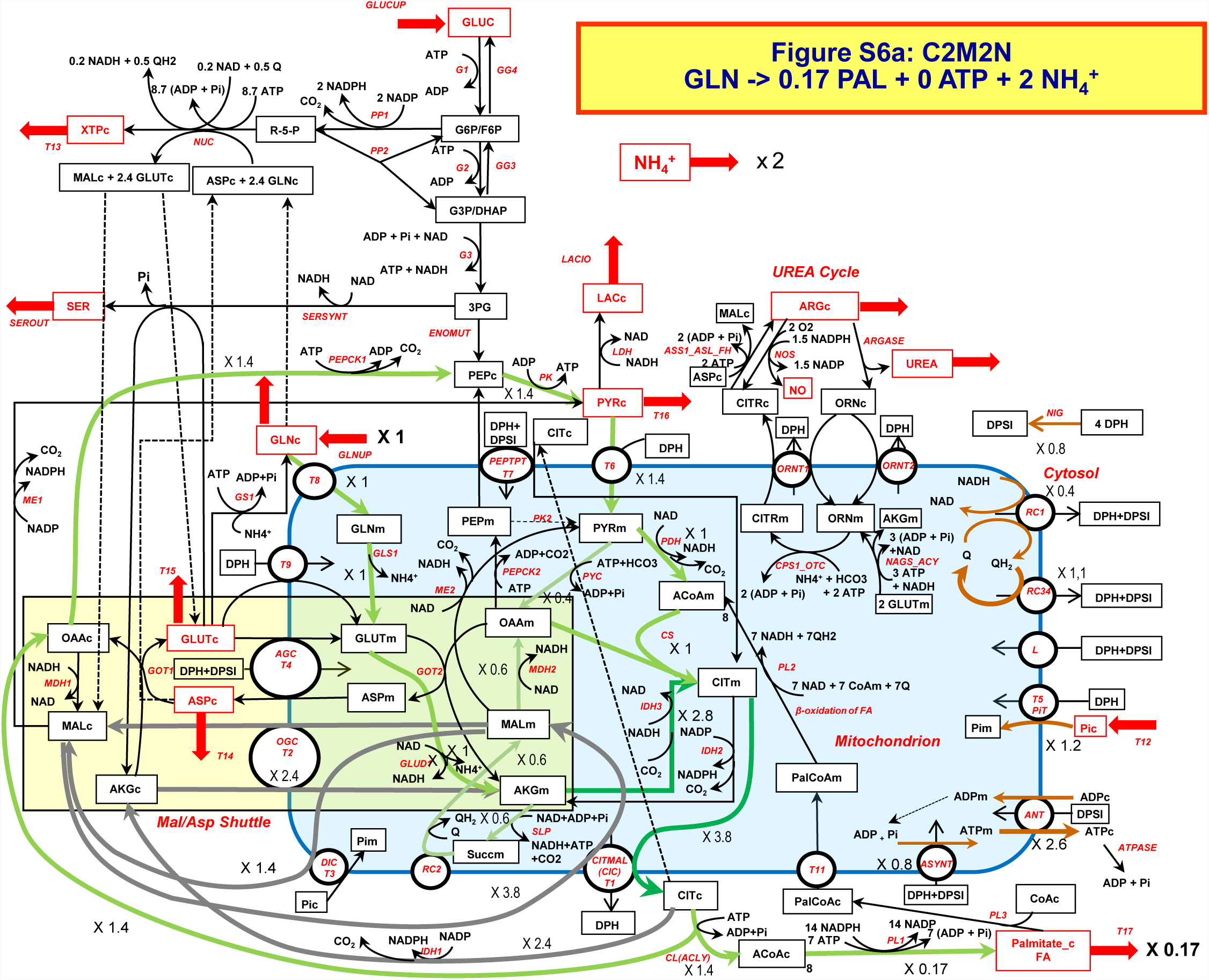
Palmitate synthesis from glutamine. Green and grey arrows represent the main pathway. The glutamine-derived AKG is split in a reductive pathway (reversion of IDH3 in dark green) and an oxidative pathway (light green). There is a cycle involving –IDH3 (dark green) and IDH1 (grey) to transform NADHm in NADPHc for fatty acids synthesis. Brown arrows depict OXPHOS.

**Figure S6b.**
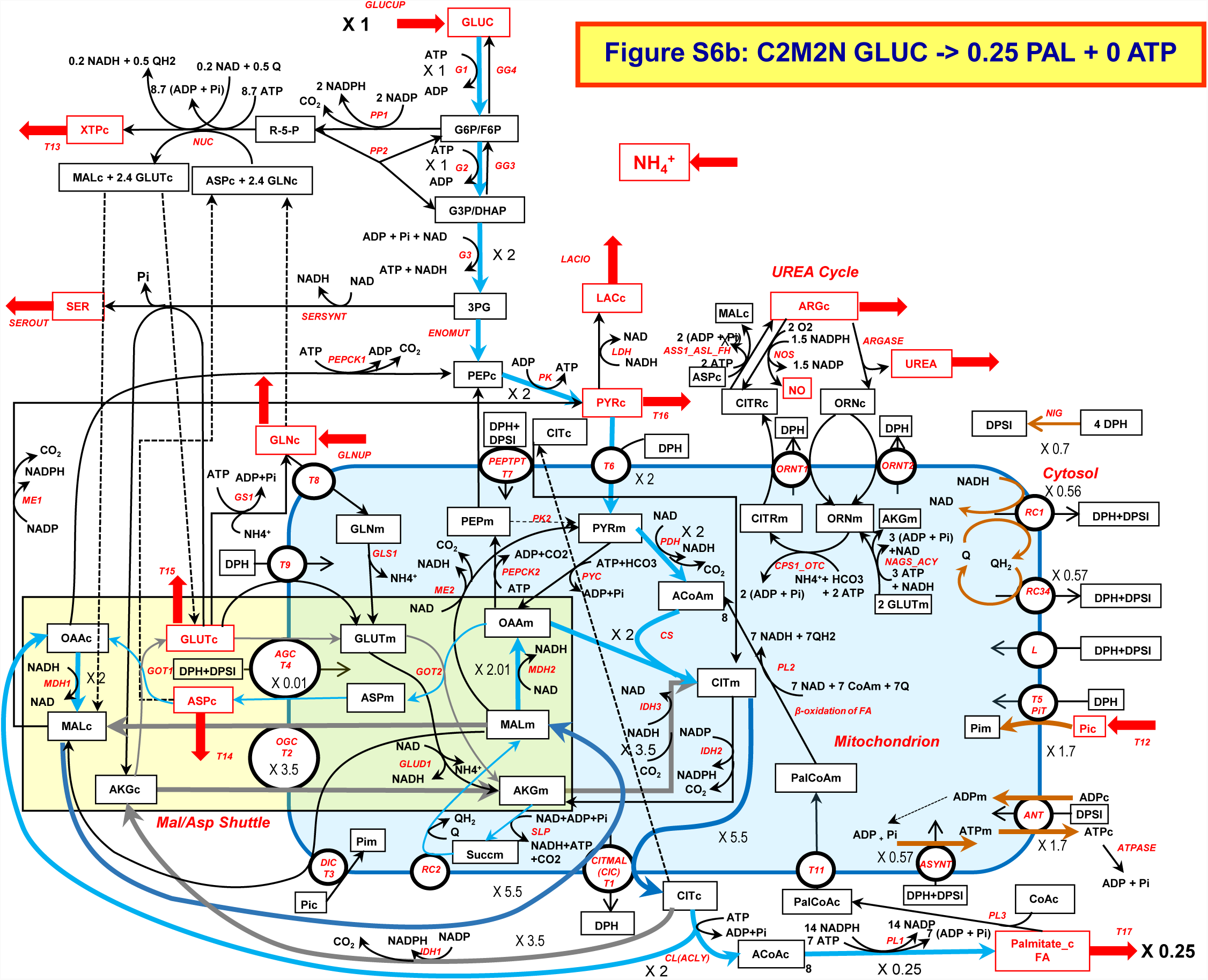
Palmitate synthesis from glucose. Blue and grey arrows represent the main pathways (flux = 2). There is a cycle of MAL (grey and dark blue) to export citrate and a cycle involving (-IDH3) and IDH1 (in grey) to transform NADHm in NADPHc necessary for fatty acids synthesis. Brown arrows depict OXPHOS.

**Figure S7a.**
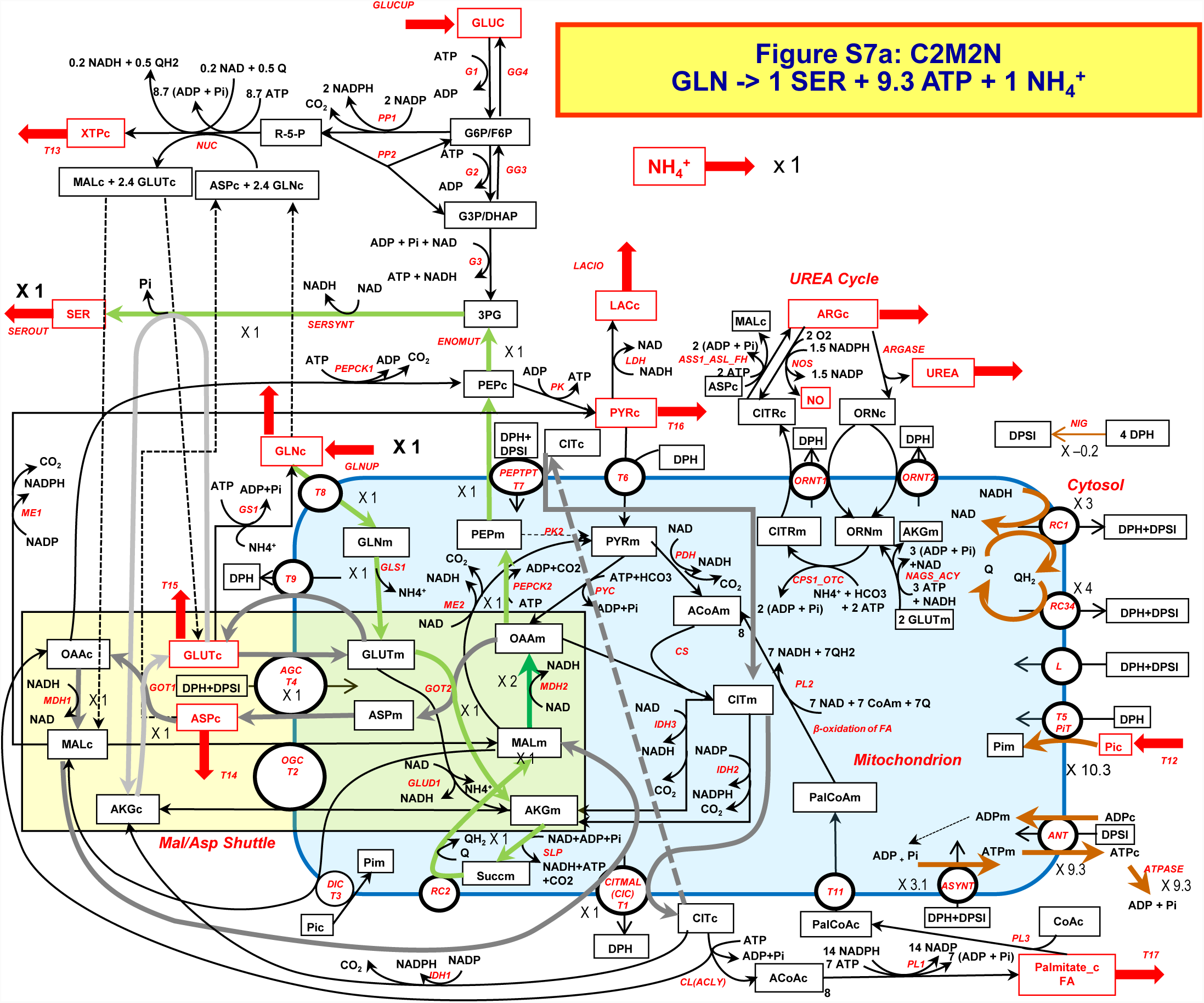
Serine synthesis from glutamine. Green and grey arrows represent the main pathways (flux = 1). The NADHc produced in serine synthesis is reoxidized by the respiratory chain through a shuttle involving the ASP/GLUT carrier T4 the CIT/MAL carrier (T1) and the T7 antiporter that insure the output of PEP against citrate for 3PG synthesis (in grey). Brown arrows depict OXPHOS.

**Figure S7b.**
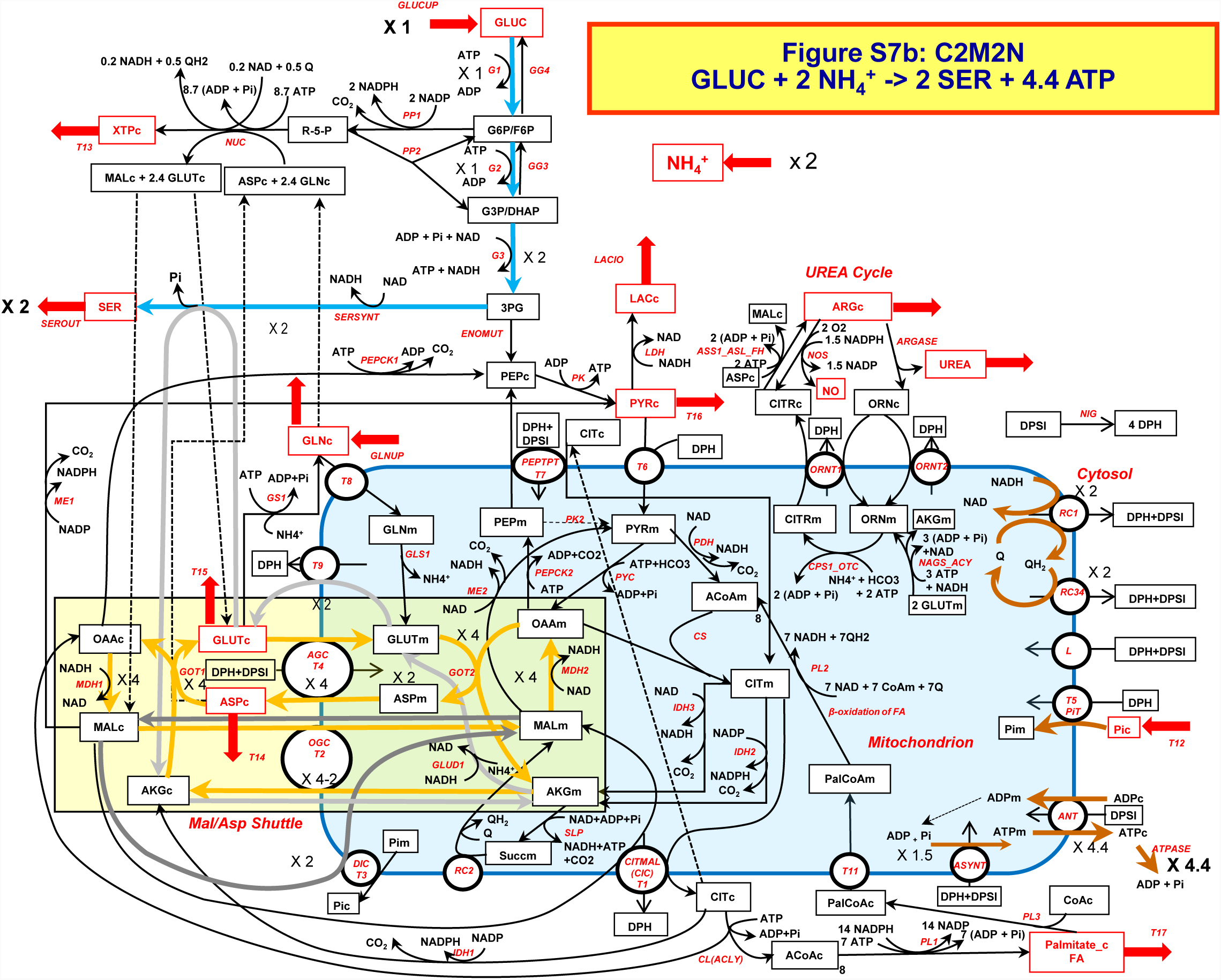
Serine synthesis from glucose. Blue and grey arrows represent the main pathway (flux = 2). Yellow arrows correspond to the malate/aspartate shuttle (flux = 4) to reoxidise NADHc produced by glycolysis and SERSYNT. Brown arrows depict OXPHOS.

**Figure S8a.**
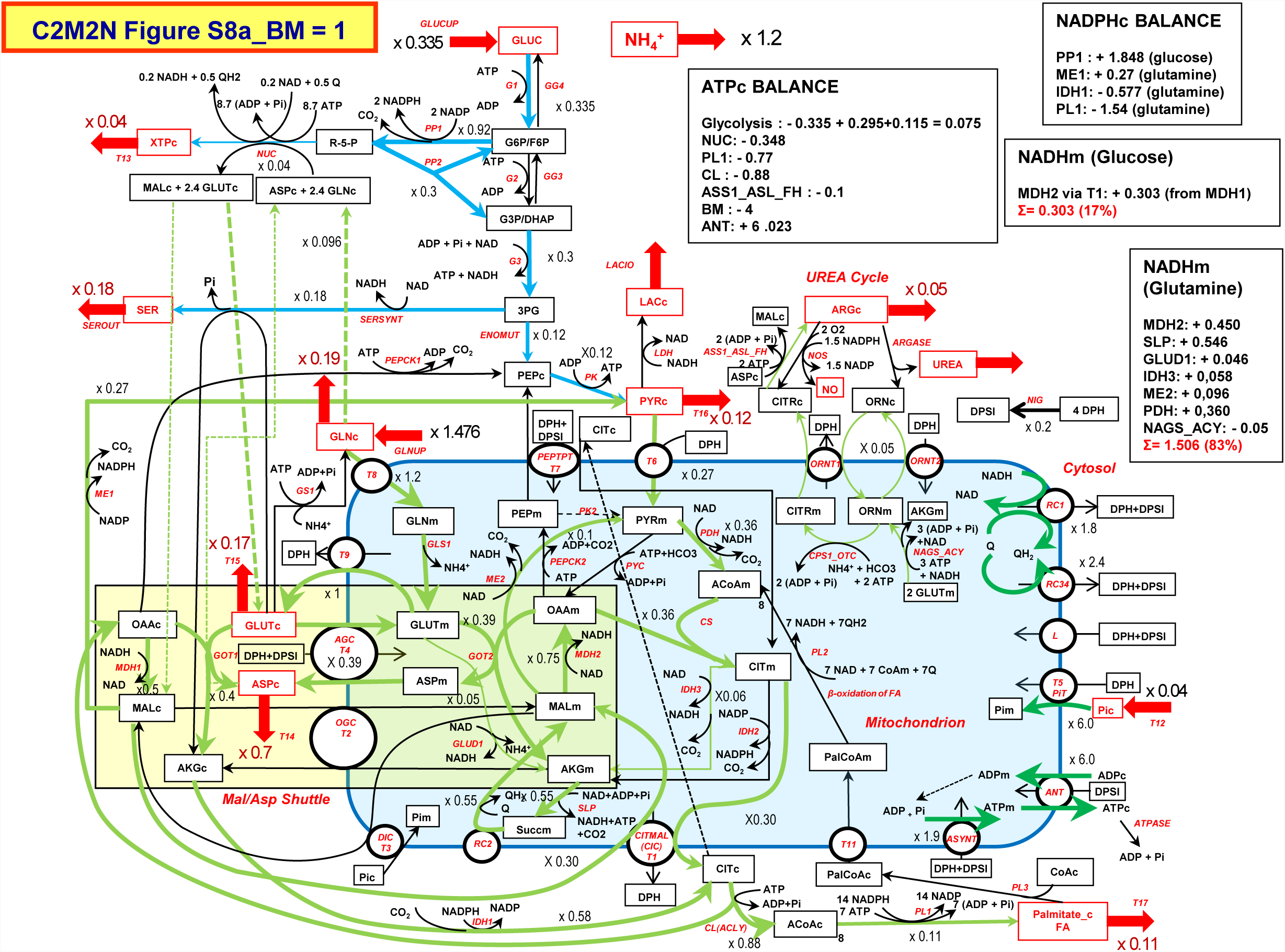
Interplay between glutamine, ammonia and glucose to sustain cell proliferation. The uptake of glucose and glutamine are free to make 1 arbitrary unit of biomass flux. Glutamine catabolism is in green and glucose catabolism in blue. Nitrogen is incorporated as a large excess of glutamine with release of ammonia. The origin of NADHm (glucose or glutamine) is indicated in the boxes showing that most of mitochondrial ATP synthesis is due to glutamine (it is the reason for dark green respiratory chain). Few ATPc is due to glycolysis (see ATPc balance box). Most of NADP/NADPH cycling is also due to glutamine.

**Figure S8b.**
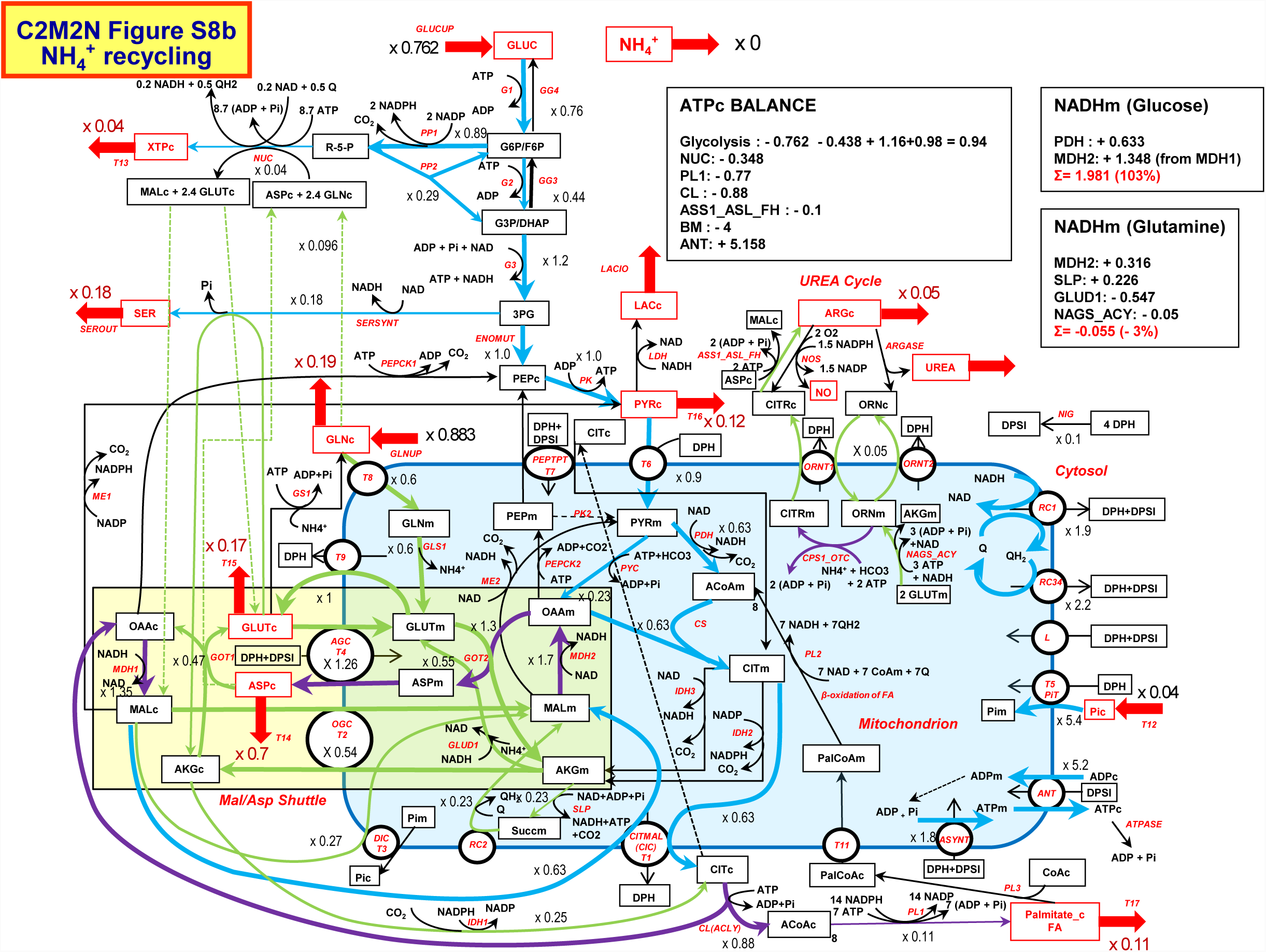
Interplay between glutamine, ammonia and glucose to sustain cell proliferation. This solution depicts a case where ammonia release is cancelled. Ammonia produced by GLS1 is thus stoichiometrically recycled mainly by GLUD1 (Glutamate dehydrogenase) in the reverse direction and to a lower extent by carbamyl phosphate synthesis (CPS1 in dark violet) to ultimately give arginine. In this simulation, nearly 100% of ATP is due to glucose (see NADHm and ATP boxes).

**Figure S8c.**
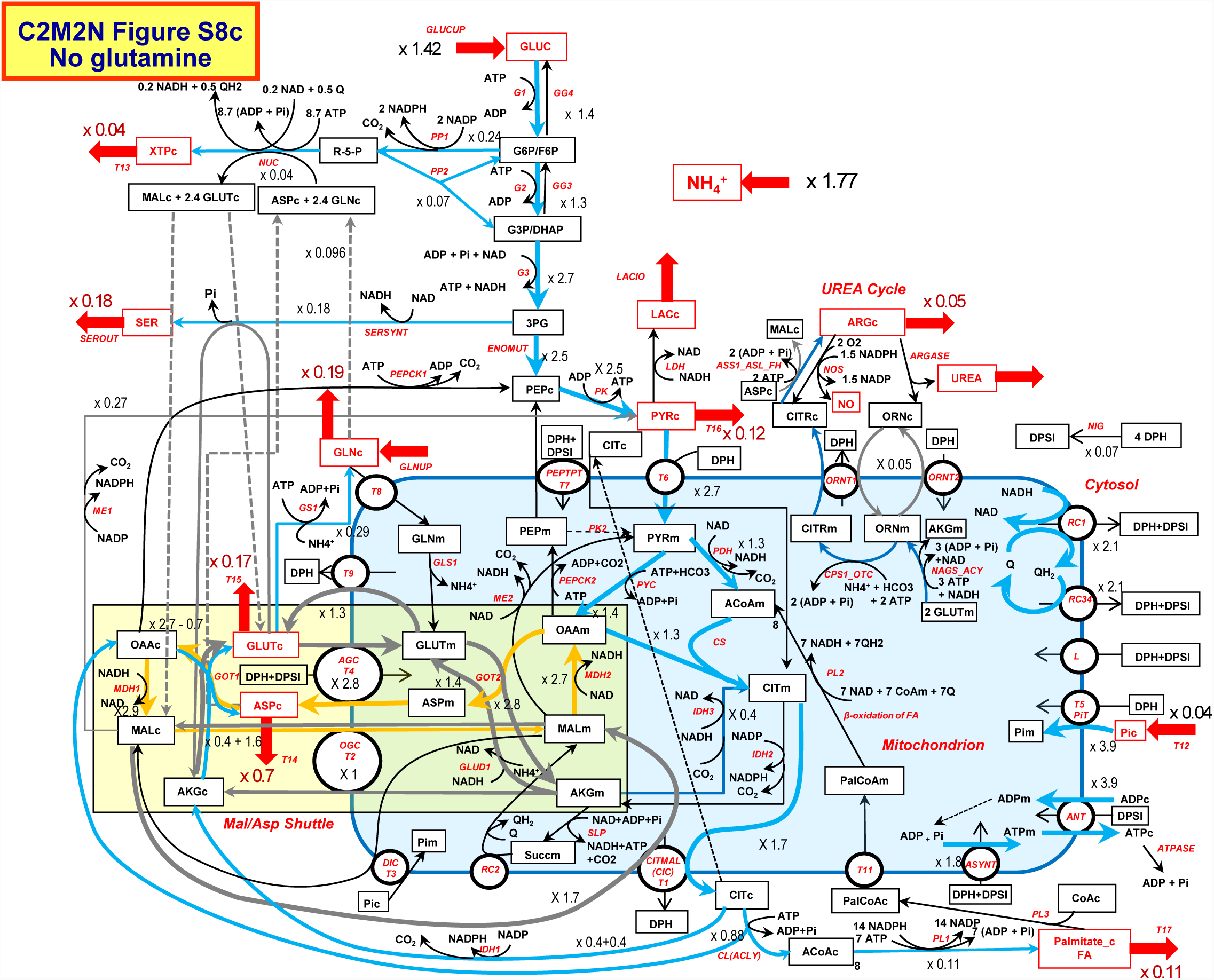
Interplay between ammonia and glucose to sustain cell proliferation. Nitrogen is incorporated as ammonia in glutamate by the reversion of glutamate dehydrogenase (-Glud1) and in glutamine by glutamine synthase (GS1). A small part of NH_4^+^_ is incorporated directly by CPS1 (carbamyl phosphate synthase) for arginine synthesis. Four cycles in yellow (MAS flux = 1.27) and in grey are operating. They are described in Supplementary Materials 2.

**Figure S9.**
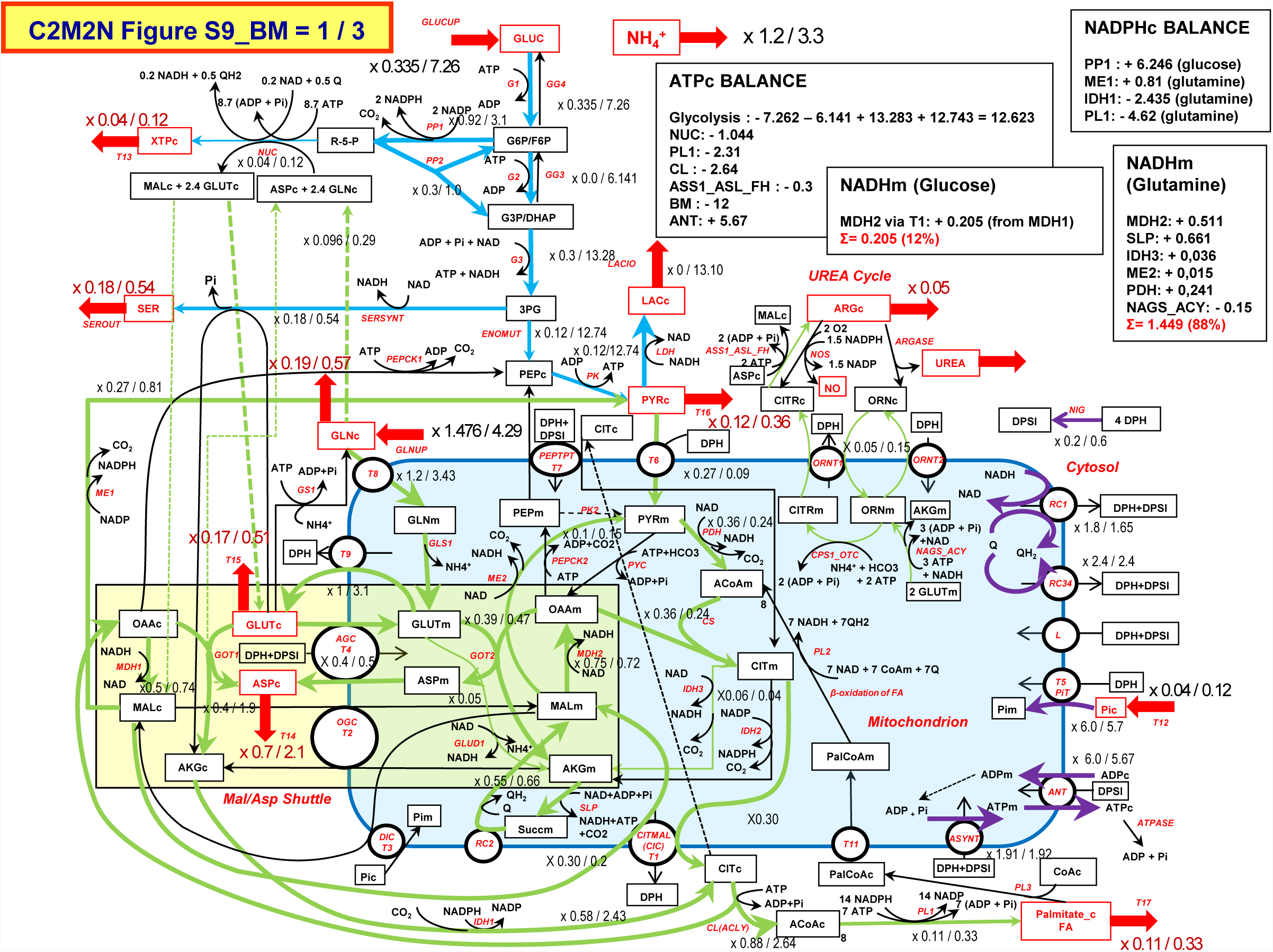
Interplay between glutamine, glucose and ammonia to sustain a flux of biomass of 1 or 3 (1/3 on the Figure). The Warburg effect. The fluxes for BM = 1 are identical to the fluxes in Figure S8a. They are kept here for comparison with the fluxes for BM = 3 in order to follow the changes of fluxes in the Warburg effect. In comparison with Figure S8a, the ANT output of mitochondrial ATP is nearly the same (see the ATPc box). The majority of ATPc comes from glycolysis with a large increase in glucose uptake accompanied by a large release of lactate. There is also a nearly 3-fold increase in glutamine uptake corresponding to the 3 fold demand in biomass. The Balance boxes are given for the case of a biomass BM = 3

## References

1. Orth, J.D.; Thiele, I.; Palsson, B.Ø. What is flux balance analysis? Nat. Biotechnol. 2010, 28, 245–248.

2. Swainston, N.; Smallbone, K.; Hefzi, H.; Dobson, P.D.; Brewer, J.; Hanscho, M.; Zielinski, D.C.; Ang, K.S.; Gardiner, N.J.; Gutierrez, J.M.; et al. Recon 2.2: from reconstruction to model of human metabolism. Metabolomics Off. J. Metabolomic Soc. 2016, 12, 109.

3. Duarte, N.C.; Becker, S.A.; Jamshidi, N.; Thiele, I.; Mo, M.L.; Vo, T.D.; Srivas, R.; Palsson, B.Ø. Global reconstruction of the human metabolic network based on genomic and bibliomic data. Proc. Natl. Acad. Sci. U. S. A. 2007, 104, 1777–1782.

4. Thiele, I.; Swainston, N.; Fleming, R.M.T.; Hoppe, A.; Sahoo, S.; Aurich, M.K.; Haraldsdottir, H.; Mo, M.L.; Rolfsson, O.; Stobbe, M.D.; et al. A community-driven global reconstruction of human metabolism. Nat. Biotechnol. 2013, 31, 419–425.

5. Ma, H.; Sorokin, A.; Mazein, A.; Selkov, A.; Selkov, E.; Demin, O.; Goryanin, I. The Edinburgh human metabolic network reconstruction and its functional analysis. Mol. Syst. Biol. 2007, 3, 135.

6. Mardinoglu, A.; Agren, R.; Kampf, C.; Asplund, A.; Nookaew, I.; Jacobson, P.; Walley, A.J.; Froguel, P.; Carlsson, L.M.; Uhlen, M.; et al. Integration of clinical data with a genome-scale metabolic model of the human adipocyte. Mol. Syst. Biol. 2013, 9, 649.

7. Mardinoglu, A.; Agren, R.; Kampf, C.; Asplund, A.; Uhlen, M.; Nielsen, J. Genome-scale metabolic modelling of hepatocytes reveals serine deficiency in patients with non-alcoholic fatty liver disease. Nat. Commun. 2014, 5, 3083.

8. Uhlén, M.; Fagerberg, L.; Hallström, B.M.; Lindskog, C.; Oksvold, P.; Mardinoglu, A.; Sivertsson, Å.; Kampf, C.; Sjöstedt, E.; Asplund, A.; et al. Proteomics. Tissue-based map of the human proteome. Science 2015, 347, 1260419.

9. Yizhak, K.; Chaneton, B.; Gottlieb, E.; Ruppin, E. Modeling cancer metabolism on a genome scale. Mol. Syst. Biol. 2015, 11, 817.

10. Bordbar, A.; Monk, J.M.; King, Z.A.; Palsson, B.O. Constraint-based models predict metabolic and associated cellular functions. Nat. Rev. Genet. 2014, 15, 107–120.

11. Smith, A.C.; Eyassu, F.; Mazat, J.-P.; Robinson, A.J. MitoCore: a curated constraint-based model for simulating human central metabolism. BMC Syst. Biol. 2017, 11, 114.

12. Mitchell, P. Coupling of phosphorylation to electron and hydrogen transfer by a chemi-osmotic type of mechanism. Nature 1961, 191, 144–148.

13. Schuster, S.; Dandekar, T.; Fell, D.A. Detection of elementary flux modes in biochemical networks: a promising tool for pathway analysis and metabolic engineering. Trends Biotechnol. 1999, 17, 53–60.

14. Metallo, C.M.; Gameiro, P.A.; Bell, E.L.; Mattaini, K.R.; Yang, J.; Hiller, K.; Jewell, C.M.; Johnson, Z.R.; Irvine, D.J.; Guarente, L.; et al. Reductive glutamine metabolism by IDH1 mediates lipogenesis under hypoxia. Nature 2011, 481, 380–384.

15. Mullen, A.R.; Wheaton, W.W.; Jin, E.S.; Chen, P.-H.; Sullivan, L.B.; Cheng, T.; Yang, Y.; Linehan, W.M.; Chandel, N.S.; DeBerardinis, R.J. Reductive carboxylation supports growth in tumour cells with defective mitochondria. Nature 2011, 481, 385–388.

16. Cluntun, A.A.; Lukey, M.J.; Cerione, R.A.; Locasale, J.W. Glutamine Metabolism in Cancer: Understanding the Heterogeneity. Trends Cancer 2017, 3, 169–180.

17. Eagle, H. The specific amino acid requirements of a human carcinoma cell (Stain HeLa) in tissue culture. J. Exp. Med. 1955, 102, 37–48.

18. Kvamme, E.; Svenneby, G. Effect of anaerobiosis and addition of keto acids on glutamine utilization by Ehrlich ascites-tumor cells. Biochim. Biophys. Acta 1960, 42, 187–188.

19. Daye, D.; Wellen, K.E. Metabolic reprogramming in cancer: unraveling the role of glutamine in tumorigenesis. Semin. Cell Dev. Biol. 2012, 23, 362–369.

20. Altman, B.J.; Stine, Z.E.; Dang, C.V. From Krebs to clinic: glutamine metabolism to cancer therapy. Nat. Rev. Cancer 2016, 16, 773.

21. Fiermonte, G.; Palmieri, L.; Todisco, S.; Agrimi, G.; Palmieri, F.; Walker, J.E. Identification of the mitochondrial glutamate transporter. Bacterial expression, reconstitution, functional characterization, and tissue distribution of two human isoforms. J. Biol. Chem. 2002, 277, 19289–19294.

22. Chen, Q.; Kirk, K.; Shurubor, Y.I.; Zhao, D.; Arreguin, A.J.; Shahi, I.; Valsecchi, F.; Primiano, G.; Calder, E.L.; Carelli, V.; et al. Rewiring of Glutamine Metabolism Is a Bioenergetic Adaptation of Human Cells with Mitochondrial DNA Mutations. Cell Metab. 2018, 27, 1007–1025.e5.

23. Boele, J.; Olivier, B.G.; Teusink, B. FAME, the Flux Analysis and Modeling Environment. BMC Syst. Biol. 2012, 6, 8.

24. Folger, O.; Jerby, L.; Frezza, C.; Gottlieb, E.; Ruppin, E.; Shlomi, T. Predicting selective drug targets in cancer through metabolic networks. Mol. Syst. Biol. 2011, 7, 501.

25. Yang, C.; Ko, B.; Hensley, C.T.; Jiang, L.; Wasti, A.T.; Kim, J.; Sudderth, J.; Calvaruso, M.A.; Lumata, L.; Mitsche, M.; et al. Glutamine oxidation maintains the TCA cycle and cell survival during impaired mitochondrial pyruvate transport. Mol. Cell 2014, 56, 414–424.

26. Fan, J.; Kamphorst, J.J.; Mathew, R.; Chung, M.K.; White, E.; Shlomi, T.; Rabinowitz, J.D. Glutamine-driven oxidative phosphorylation is a major ATP source in transformed mammalian cells in both normoxia and hypoxia. Mol. Syst. Biol. 2013, 9, 712.

27. Birsoy, K.; Wang, T.; Chen, W.W.; Freinkman, E.; Abu-Remaileh, M.; Sabatini, D.M. An Essential Role of the Mitochondrial Electron Transport Chain in Cell Proliferation Is to Enable Aspartate Synthesis. Cell 2015, 162, 540–551.

28. Van Vranken, J.G.; Rutter, J. You Down With ETC? Yeah, You Know D! Cell 2015, 162, 471–473.

29. Sullivan, L.B.; Gui, D.Y.; Hosios, A.M.; Bush, L.N.; Freinkman, E.; Vander Heiden, M.G. Supporting Aspartate Biosynthesis Is an Essential Function of Respiration in Proliferating Cells. Cell 2015, 162, 552–563.

30. Garcia-Bermudez, J.; Baudrier, L.; La, K.; Zhu, X.G.; Fidelin, J.; Sviderskiy, V.O.; Papagiannakopoulos, T.; Molina, H.; Snuderl, M.; Lewis, C.A.; et al. Aspartate is a limiting metabolite for cancer cell proliferation under hypoxia and in tumours. Nat. Cell Biol. 2018, 20, 775.

31. Sullivan, L.B.; Luengo, A.; Danai, L.V.; Bush, L.N.; Diehl, F.F.; Hosios, A.M.; Lau, A.N.; Elmiligy, S.; Malstrom, S.; Lewis, C.A.; et al. Aspartate is an endogenous metabolic limitation for tumour growth. Nat. Cell Biol. 2018, 20, 782.

32. DeBerardinis, R.J.; Mancuso, A.; Daikhin, E.; Nissim, I.; Yudkoff, M.; Wehrli, S.; Thompson, C.B. Beyond aerobic glycolysis: transformed cells can engage in glutamine metabolism that exceeds the requirement for protein and nucleotide synthesis. Proc. Natl. Acad. Sci. U. S. A. 2007, 104, 19345–19350.

33. Wise, D.R.; Ward, P.S.; Shay, J.E.S.; Cross, J.R.; Gruber, J.J.; Sachdeva, U.M.; Platt, J.M.; DeMatteo, R.G.; Simon, M.C.; Thompson, C.B. Hypoxia promotes isocitrate dehydrogenase-dependent carboxylation of α-ketoglutarate to citrate to support cell growth and viability. Proc. Natl. Acad. Sci. U. S. A. 2011, 108, 19611–19616.

34. Gaude, E.; Schmidt, C.; Gammage, P.A.; Dugourd, A.; Blacker, T.; Chew, S.P.; Saez-Rodriguez, J.; O’Neill, J.S.; Szabadkai, G.; Minczuk, M.; et al. NADH Shuttling Couples Cytosolic Reductive Carboxylation of Glutamine with Glycolysis in Cells with Mitochondrial Dysfunction. Mol. Cell 2018, 69, 581–593.e7.

35. Fendt, S.-M.; Bell, E.L.; Keibler, M.A.; Olenchock, B.A.; Mayers, J.R.; Wasylenko, T.M.; Vokes, N.I.; Guarente, L.; Vander Heiden, M.G.; Stephanopoulos, G. Reductive glutamine metabolism is a function of the α-ketoglutarate to citrate ratio in cells. Nat. Commun. 2013, 4, 2236.

36. Corbet, C.; Draoui, N.; Polet, F.; Pinto, A.; Drozak, X.; Riant, O.; Feron, O. The SIRT1/HIF2α axis drives reductive glutamine metabolism under chronic acidosis and alters tumor response to therapy. Cancer Res. 2014, 74, 5507–5519.

37. Corbet, C.; Feron, O. Metabolic and mind shifts: from glucose to glutamine and acetate addictions in cancer. Curr. Opin. Clin. Nutr. Metab. Care 2015, 18, 346–353.

38. Hanse, E.A.; Ruan, C.; Kachman, M.; Wang, D.; Lowman, X.H.; Kelekar, A. Cytosolic malate dehydrogenase activity helps support glycolysis in actively proliferating cells and cancer. Oncogene 2017, 36, 3915–3924.

39. Kalhan, S.C.; Hanson, R.W. Resurgence of serine: an often neglected but indispensable amino Acid. J. Biol. Chem. 2012, 287, 19786–19791.

40. Snell, K. Enzymes of serine metabolism in normal, developing and neoplastic rat tissues. Adv. Enzyme Regul. 1984, 22, 325–400.

41. Possemato, R.; Marks, K.M.; Shaul, Y.D.; Pacold, M.E.; Kim, D.; Birsoy, K.; Sethumadhavan, S.; Woo, H.-K.; Jang, H.G.; Jha, A.K.; et al. Functional genomics reveal that the serine synthesis pathway is essential in breast cancer. Nature 2011, 476, 346–350.

42. Locasale, J.W.; Grassian, A.R.; Melman, T.; Lyssiotis, C.A.; Mattaini, K.R.; Bass, A.J.; Heffron, G.; Metallo, C.M.; Muranen, T.; Sharfi, H.; et al. Phosphoglycerate dehydrogenase diverts glycolytic flux and contributes to oncogenesis. Nat. Genet. 2011, 43, 869–874.

43. Pollari, S.; Käkönen, S.-M.; Edgren, H.; Wolf, M.; Kohonen, P.; Sara, H.; Guise, T.; Nees, M.; Kallioniemi, O. Enhanced serine production by bone metastatic breast cancer cells stimulates osteoclastogenesis. Breast Cancer Res. Treat. 2011, 125, 421–430.

44. Beauvoit, B.; Colombié, S.; Issa, R.; Mazat, J.-P.; Nazaret, C.; Pérès, S. Human-Scale Metabolic Network of Central Carbon Metabolism. Application to serine metabolism from glutamine in Cancer Cells. In Proceedings of the Évry Spring school on advances in systems and synthetic biology, March 21th-25th, 2016; Amar, P., Képès, F., Norris, V., Eds.; 2016; pp. 37–56.

45. Kalhan, S.C.; Uppal, S.O.; Moorman, J.L.; Bennett, C.; Gruca, L.L.; Parimi, P.S.; Dasarathy, S.; Serre, D.; Hanson, R.W. Metabolic and genomic response to dietary isocaloric protein restriction in the rat. J. Biol. Chem. 2011, 286, 5266–5277.

46. Takeuchi, Y.; Nakayama, Y.; Fukusaki, E.; Irino, Y. Glutamate production from ammonia via glutamate dehydrogenase 2 activity supports cancer cell proliferation under glutamine depletion. Biochem. Biophys. Res. Commun. 2018, 495, 761–767.

47. Spinelli, J.B.; Yoon, H.; Ringel, A.E.; Jeanfavre, S.; Clish, C.B.; Haigis, M.C. Metabolic recycling of ammonia via glutamate dehydrogenase supports breast cancer biomass. Science 2017, 358, 941–946.

48. Eng, C.H.; Yu, K.; Lucas, J.; White, E.; Abraham, R.T. Ammonia derived from glutaminolysis is a diffusible regulator of autophagy. Sci. Signal. 2010, 3, ra31.

49. Dang, C.V. Feeding frenzy for cancer cells. Science 2017, 358, 862–863.

50. Martinez-Outschoorn, U.E.; Peiris-Pagés, M.; Pestell, R.G.; Sotgia, F.; Lisanti, M.P. Cancer metabolism: a therapeutic perspective. Nat. Rev. Clin. Oncol. 2017, 14, 11–31.

51. Süer Gökmen, S.; Yörük, Y.; Cakir, E.; Yorulmaz, F.; Gülen, S. Arginase and ornithine, as markers in human non-small cell lung carcinoma. Cancer Biochem. Biophys. 1999, 17, 125–131.

52. Singh, R.; Pervin, S.; Karimi, A.; Cederbaum, S.; Chaudhuri, G. Arginase activity in human breast cancer cell lines: N(omega)-hydroxy-L-arginine selectively inhibits cell proliferation and induces apoptosis in MDA-MB-468 cells. Cancer Res. 2000, 60, 3305–3312.

53. Keshet, R.; Szlosarek, P.; Carracedo, A.; Erez, A. Rewiring urea cycle metabolism in cancer to support anabolism. Nat. Rev. Cancer 2018, 18, 634.

54. Long, Y.; Tsai, W.-B.; Wangpaichitr, M.; Tsukamoto, T.; Savaraj, N.; Feun, L.G.; Kuo, M.T. Arginine deiminase resistance in melanoma cells is associated with metabolic reprogramming, glucose dependence, and glutamine addiction. Mol. Cancer Ther. 2013, 12, 2581–2590.

55. Dai, Z.; Shestov, A.A.; Lai, L.; Locasale, J.W. A Flux Balance of Glucose Metabolism Clarifies the Requirements of the Warburg Effect. Biophys. J. 2016, 111, 1088–1100.

56. Gellerich, F.N.; Gizatullina, Z.; Gainutdinov, T.; Muth, K.; Seppet, E.; Orynbayeva, Z.; Vielhaber, S. The control of brain mitochondrial energization by cytosolic calcium: the mitochondrial gas pedal. IUBMB Life 2013, 65, 180–190.

